# *Dppa2/4* Counteract *De Novo* Methylation to Establish a Permissive Epigenome for Development

**DOI:** 10.1101/2020.03.11.987701

**Authors:** Kristjan H. Gretarsson, Jamie A. Hackett

## Abstract

Early mammalian development entails genome-wide epigenome remodeling, including DNA methylation erasure and reacquisition, which facilitates developmental competence. To uncover the mechanisms that orchestrate DNA methylation (DNAme) dynamics, we coupled a single-cell ratiometric DNAme reporter with unbiased CRISPR screening in ESC. We identify key genes and regulatory pathways that drive global DNA hypomethylation, and characterise roles for *Cop1* and *Dusp6*. We also identify *Dppa2* and *Dppa4* as essential safeguards of focal epigenetic states. In their absence, developmental genes and evolutionary-young LINE1 elements, which DPPA2 specifically binds, lose H3K4me3 and gain ectopic *de novo* DNA methylation in pluripotent cells. Consequently, lineage-associated genes (and LINE1) acquire a repressive epigenetic memory, which renders them incompetent for activation during future lineage-specification. *Dppa2/4* thereby sculpt the pluripotent epigenome by facilitating H3K4me3 and bivalency to counteract *de novo* methylation; a function co-opted by evolutionary young LINE1 to evade epigenetic decommissioning.

## INTRODUCTION

Mammalian fertilisation is accompanied by widespread epigenetic remodeling of inherited genomes, including global DNA demethylation and reorganization of chromatin landscapes^1–4^. This epigenetic resetting equalises the distinct parental epigenomes, and also correlates with the emergence of naïve pluripotency, implying epigenome remodeling is central to establish developmental competence. Such ‘competence’ confers the capacity of the genome to transcriptionally respond to future inductive signals for multiple lineages, and can be considered the execution of pluripotency. This is particularly critical for lineage-associated genes that need to be transiently repressed during pluripotent phases, whilst remaining competent (primed) for robust activation in subsets of forthcoming cell fates^5^. Indeed, the importance of a permissive epigenome is supported by observations of impaired or reduced developmental competence after somatic cell nuclear transfer (SCNT) or in induced pluripotent stem (iPS) cells, which are susceptible to incomplete epigenetic resetting^6,7^. Investigating the complex mechanisms that underpin epigenome (re)programming is therefore an important focus towards understanding developmental potency.

Several lines of evidence indicate that resetting DNA methylation (DNAme) during development is mediated by parallel mechanisms^8^. Amongst these, repression of the maintenance DNA methylation machinery is central and appears to occur through post-translational regulation of UHRF1^9,10^, at least in part via STELLA activity^11,12^. This is further supported by PRDM14, which suppresses the *de novo* methylases, and is necessary for DNA hypomethylation in naïve pluripotent cells^13,14^. In parallel, replication-independent DNAme erasure occurs on both the maternal and paternal genomes^1^. Counterintuitively, *de novo* methylation remains active throughout epigenetic reprogramming but is offset, in part, via TET proteins^15^. These collective mechanisms contribute towards resetting the epigenome, but also present an opportunity for transposable elements (TE), such as LINE1, to mobilise due to epigenetic de-restriction. Such LINE1 activation has been linked with key developmental events^16^, but could also represent a hazard to the genome if left unrestrained^17,18^. Epigenetic (re)programming therefore likely strikes a balance between genome-wide resetting to a competent state for development, and targeted regulation. Nevertheless, a complete understanding of the mechanisms that cross-talk to remodel the epigenome, how they interact to balance focal and global effects, and what the full repertoire of genes involved is lacking.

Here we have coupled a single-cell ratiometric reporter of cellular DNA methylation status with CRISPR screening to unbiasedly identify the gene networks that underpin DNAme remodeling. In doing so we identify upstream regulators of global DNAme erasure in pluripotent cells. We also identify *Dppa2* and *Dppa4* as key genes that safeguard against focal *de novo* DNA methylation and epigenetic silencing at lineage-associated genes by integrating chromatin states, and consequently confer developmental competence. Remarkably, LINE1 elements appear to have exapted this *Dppa2/4* function to escape epigenetic surveillance and enable competence for precocious activation, potentially highlighting an evolving genomic conflict.

## RESULTS

### Single-cell monitoring of DNA demethylation

To identify regulators of epigenetic remodeling we exploited the reporter for genomic DNA <u>m</u>ethylation (RGM)^19^, which tracks the dynamic DNA methylation state of single-cells with GFP. We optimised the system for CRISPR screening in two ways. First, we replaced the original *Snrpn* imprinted promoter for the core *Kcnq1ot1* imprinted promoter, which <u>e</u>nhanced the dynamic range of reporter activity (*<u>e</u>*RGM), enabling better separation of hypo− and hyper-methylated cells (Fig S1A). Second, we converted the read-out to a ratiometric measure by introducing an additional *Ef1a*-mCherry that is not sensitive to DNA methylation (Fig 1A). This enables a single-cell ratiometric score (*e*RGM(GFP):*Ef1a*-mCherry) that normalises for general confounding effects on a reporter in a screen (*e.g.* disruption of translation factors) or inherent cell-cell variance (*e.g.* cell cycle stage).

**Figure 1.**
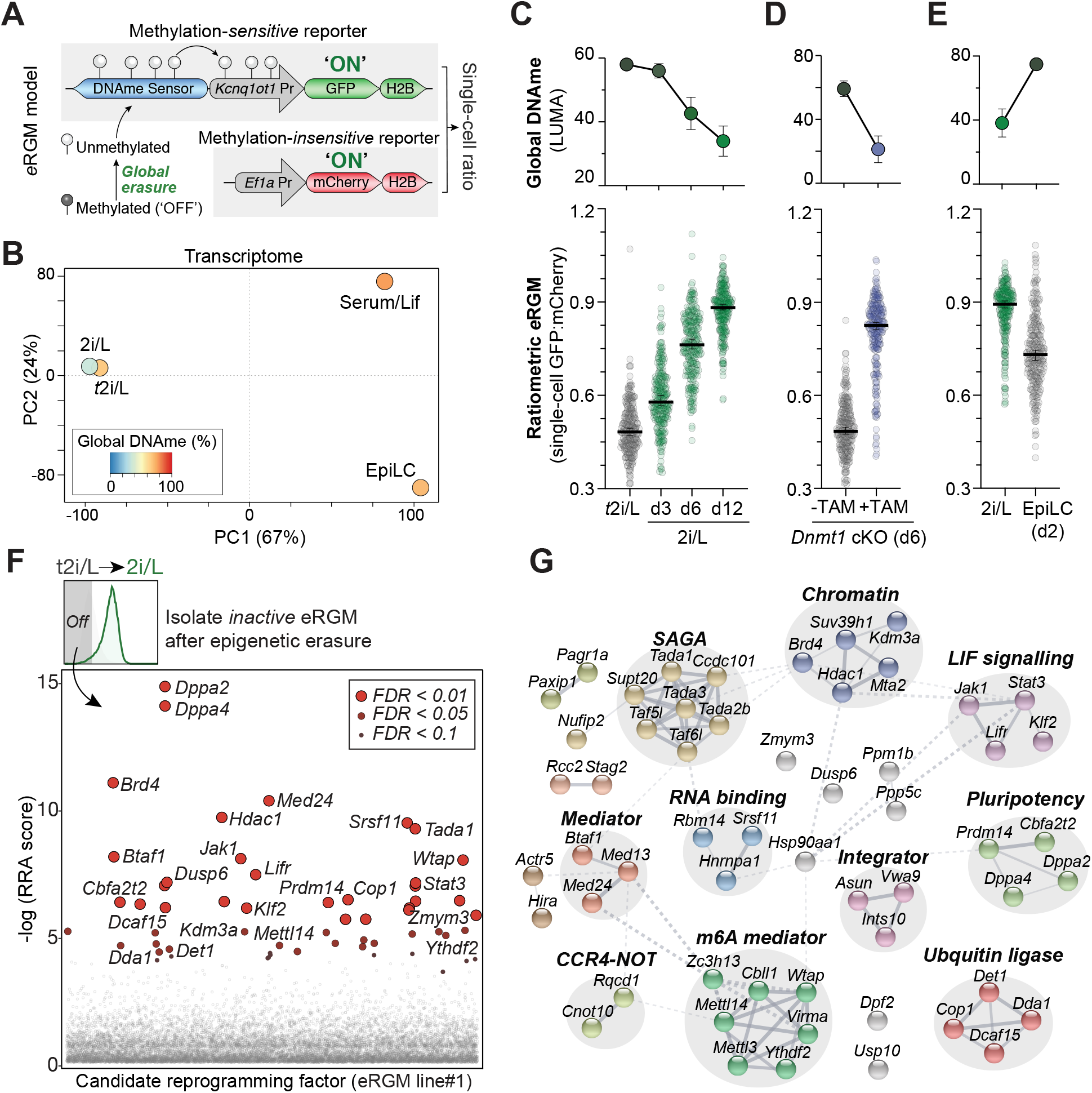
Developmental model and ratiometric reporter for DNA demethylation CRISPR screening. **A.)** Schematic of the ratiometric *e*RGM real-time DNAme reporter whereby GFP is OFF in hypermethylated cells but activates upon hypomethylation via imprinted promoter. mCherry remains active establishing a single-cell ratio score. **B.)** PCA of the transcriptomes (RNAseq) of S/L, *t*2i/L and 2i/L cultured ESC, and EpiLC, shaded by global DNA methylation level. **C-E.)** Plot showing representative single-cell (n=250) ratiometric scores of eRGM **C.)** during transition from *t*2i/L to 2i/L, **D.)** after TAM induced *Dnmt1* KO or **E.)** after EpiLC differentiation. Upper panels show corresponding changes in global DNA methylation. **F**.) Significance scores (RRA) of CRISPR knockout screen candidates required for *e*RGM (line#1) activation after *t*2i/L to 2i/L DNA demethylation transition. **G.)** STRING clustering of significant candidates from independent reprogramming screens.

To test ratiometric *e*RGM we developed a model of developmentally induced DNA demethylation. Here, murine embryonic stem cells (ESC) are maintained in a *<u>t</u>itrated* 2i/L (*t*2i/L) condition (see methods) to promote high global levels of DNA methylation (range: 64%-58%), and are then transitioned to 2i/L status to induce global demethylation (range: 30%-44%; *p*=0.0002) (Fig 1b). Importantly, global DNA demethylation after switching from *t*2i/L→2i/L occurs without significant changes in cell identity, as judged by transcriptome, which is in contrast to the switch from conventional serum/LIF to 2i/L that constitutes a major transcriptional shift^20–22^ (Fig 1B & S1B). Moreover, the induced DNA hypomethylation pattern is well correlated with developmentally imposed DNA demethylation *in vivo* (Fig S1C). Thus, the *t*2i/L→2i/L model specifically captures an authentic global epigenetic transition, including global DNA demethylation, without changes in cell identity that could confound a screen for epigenome regulators.

We next examined the capacity to detect DNA demethylation events in single-cells by generating independent ESC lines carrying the ratiometric *e*RGM system. In *t*2i/L *e*RGM was silenced in >95% of cells, consistent with high global DNA methylation. In contrast, *e*RGM exhibited a progressive activation concomitant with induced DNA methylation erasure in 2i/L, leading to *e*RGM activation in 12% of single-cells after 3 days, 67% after 6 days, and in >95% of cells upon complete DNA hypomethylation at 12 days (Fig 1C & S1D). Independent *e*RGM lines exhibited consistent response to induced hypomethylation (Fig S1D). Notably, *Ef1*-mCherry did not alter expression during this transition enabling its use as a ratiometric normaliser (Fig S1D). To further confirm *e*RGM directly reports cellular DNA methylation status, we used ESC wherein tamoxifen (TAM) drives CRE-mediated deletion of *Dnmt1* (cDKO) and, consequently, global DNA demethylation occurs independent of culture condition ^23^. Upon TAM exposure we observed a strong and progressive activation of *e*RGM amongst single-cells concomitant with *Dnmt1* (cDKO)-induced DNA hypomethylation (Fig 1D).

Finally, we tested whether *e*RGM can also respond reciprocally to *acquisition* of DNA methylation by inducing differentiation of hypomethylated ESC (in 2i/L) into hypermethylated EpiLC (global 5mC 33%→75%). Here the reporter initiated rapid silencing in parallel with induction of DNA hypermethylation (Fig 1E). We conclude the enhanced ratiometric reporter of genomic DNAme (*e*RGM) represents a single-cell read out for dynamic transitions of cellular DNA methylation status.

### A CRISPR screen for regulators of dynamic DNA methylation

To identify critical factors for DNA methylation resetting, we generated independent ESC lines carrying ratiometric *e*RGM and a single-copy of *spCas9*, and introduced into them a CRISPR knockout gRNA library ^24^. To validate the capacity of the strategy to detect epigenetic regulators, we isolated ESC that *activated e*RGM under hypermethylated (*t*2i/L) conditions, which is predicted to identify factors necessary to maintain DNA methylation and/or epigenetic silencing. Analysis using MAGeCK ^25^, revealed the top hits comprised the key machinery for maintenance DNA methylation, in particular *Dnmt1* (rank: 5, FDR=*0.00049*) and *Uhrf1* (rank: 48, FDR=*0.066*), unbiasedly confirming *e*RGM sensitivity to DNA hypomethylation (Fig S1E). We also identified regulators of chromatin-mediated silencing such as *Setdb1* (rank: 51, FDR=*0.073*,) and the HUSH complex (*Mphosph8* rank: *6*; *Morc2a* rank: *9*; *Fam208a*; rank: *13*). These data support *e*RGM specificity for detecting developmental epigenome regulators, including of cellular DNA methylation status.

We next aimed to identify factors that contribute to resetting the epigenome at focal or global scales. We induced global DNA demethylation and isolated individual ESC that failed to ratiometrically activate *e*RGM, indicative of a failure to undergo epigenetic resetting (Fig 1F). Importantly, this population was highly enriched for knockout of *Prdm14* (rank: 16, FDR=*0.0006*) the key regulator known to instruct global DNA demethylation^13^ as well as its heterodimeric co-factor *Cbfa2t2a* (rank: 20, FDR=*0.0006*)^26,27^, supporting the sensitivity of the strategy for identifying reprogramming factors (Fig 1F). Moreover, screens of independent *e*RGM ESC lines identified highly correlated (*p*=0.01, spearman RRA) candidates (Fig S2A) suggesting the system is robust. We therefore intersected significant hits (FDR<*0.05*, FC>3) from independent screens to identify a core candidate list of 56 putative genes linked with resetting the epigenome (Table S1).

Gene ontology analysis of these candidates suggested they are enriched for nuclear localisation (FDR=*3.2^−16^*), with roles related to ‘nucleic acid metabolism (FDR=*1.1^−7^*) and ‘histone modification’ (FDR=*4.3^−10^*) (Fig S2B), consistent with epigenome regulation. Applying MCF in STRING revealed they assembled into 13 clusters, which are enriched for functional interactions (PPI enrichment *p*<1.0^−16^), implying candidate genes are linked into common molecular pathways (Fig 1G). Amongst these, a cluster corresponding to the LIF/JAK/STAT3 axis emerged, supporting the previously observed role for LIF signaling in promoting DNA hypomethylation in naïve ESC^22,28^. Moreover, a cluster linked with pluripotency included the most significant hits *Dppa2* and *Dppa4*, whilst another included chromatin regulators such as *Brd4* and *Kdm3a*. We also observed multiple candidate factors involved in m6a RNA methylation (*Virma*, *Ythdf2*, *Mettl14*, *Z3h13*, *Mettl3*, *Cbll1*), ubiquitin ligases (*Cop1*, *Det1*, *Dda1*, *Dcaf15*), and phosphatases (*Dusp6, Ppm1b, Ppp5c*) (Fig 1G), which could potentially indirectly influence epigenetic status through acting as upstream regulators.

To validate the CRISPR screen hits we generated knock-out (KO) ESC populations for 24 selected candidates, by introducing a constitutive gRNA into *e*RGM*:Cas9* lines via piggyBac, and transited to hypomethylated conditions. Strikingly, knockout of each candidate resulted in a degree of impaired *e*RGM activation, implying altered epigenome remodeling in their absence (Fig 2A). This effect was robust since we generated additional knockouts in an independent *e*RGM line, with similar outcomes (Fig S2C). Interestingly, the response kinetics of *e*RGM during transition to 2i/L varied amongst candidate KO. For example, *Jak1*, *Dppa2*, *Dppa4* and *Brd4* mutants failed to activate *e*RGM *per se*, indicating a general block. In contrast, other candidate knockouts such as *Dusp6*, *Kdm3a*, *Nufip1* and *Cop1* exhibited late-onset heterogeneous activation amongst single-cells, implying delayed demethylation dynamics and reduced robustness in their absence. (Fig 2B & S2D). These validations suggest that candidate factors influence both the kinetics and absolute response of *e*RGM.

**Figure 2.**
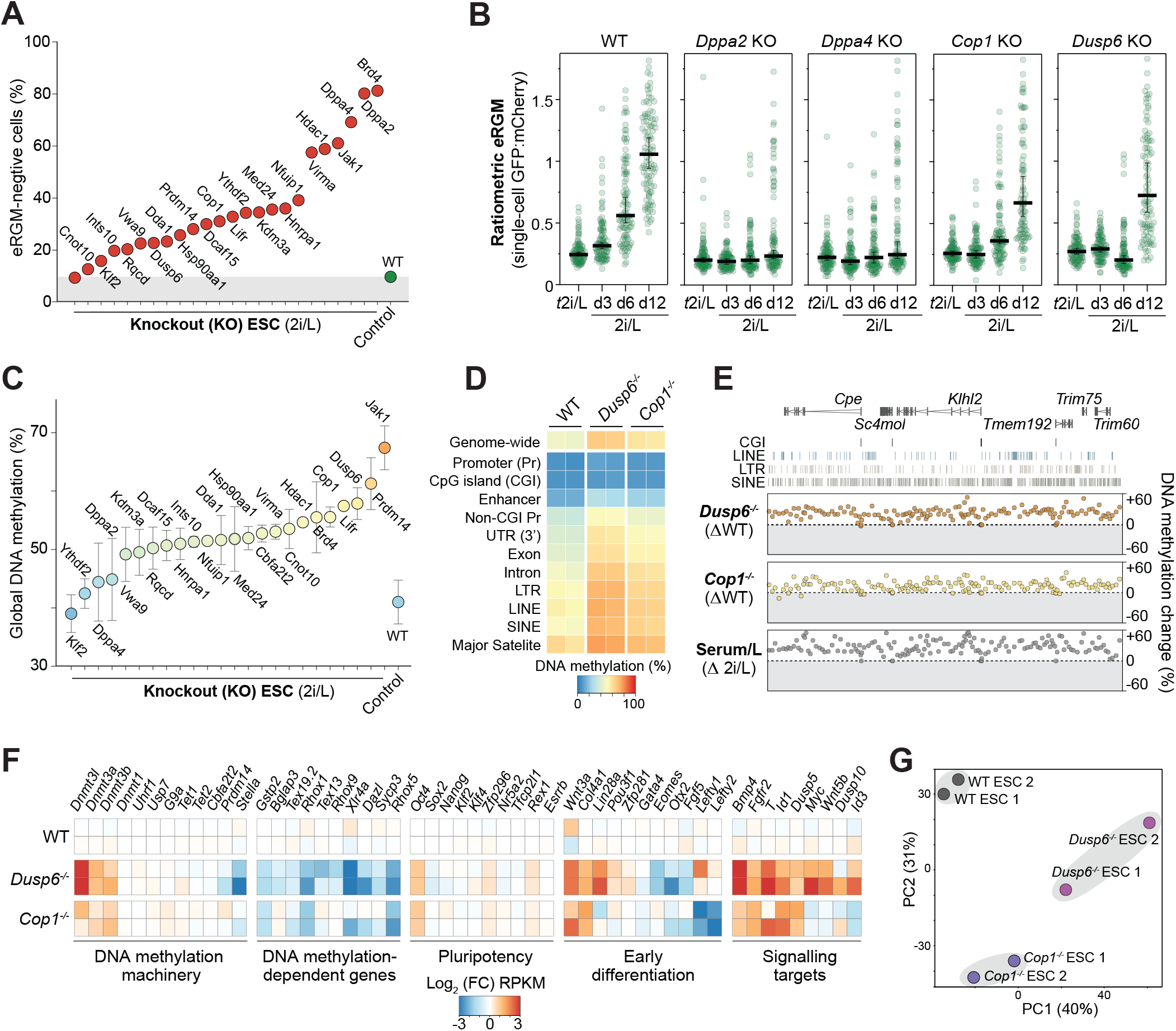
Validation of candidate DNAme reprogramming factors reveals a role for *Dusp6* and *Cop1*. **A.)** Percent of single-cells with *e*RGM ‘off’ (marked in grey) in hypomethylated 2i/L following knockout of indicated candidate genes (*n=24*) and control (non-targeting gRNA) **B.)** Dynamics of ratiometric *e*RGM activation amongst single-cells during *t*2i/L to 2i/L transition in WT, *Dppa2-/-*, *Dppa4-/-*, *Dusp6-/-* and *Cop1-/-* lines. **C.)** LUMA quantification of global DNA methylation levels in indicated knockout lines of candidate genes. **D.)** Heatmap showing DNA methylation levels genome-wide and at indicated genomic features in WT, *Dusp6-/-* and *Cop1-/-* naïve ESC. **E.)** Representative genome view of DNA hypermethylation in *Dusp6-/-* and *Cop1-/-* ESC in 2i/L, shown as a delta with WT ESC. Each datapoint represents average methylation change over a 100 CpG-tile. **F.)** Heatmap of gene expression (RNA-seq) changes for selected pathways in *Dusp6-/-* and *Cop1-/-* **G.)** Principal component analysis of global transcriptomes in WT, *Dusp6-/-* and *Cop1-/-* ESC.

To test whether the impaired *e*RGM response in candidate KO is directly indicative of incomplete epigenetic reprogramming, we used LUMA to quantitatively assess global DNA methylation. Consistent with *e*RGM, we found knockout of 20 of 24 candidate factors resulted in impaired global DNA demethylation across independent lines (Fig 2C). Amongst these is the known epigenetic regulator *Prdm14*, which maintained 64-58% global DNAme relative to hypomethylated WT control (39%), as well *Cbfa2t2* (54-52%). Novel candidates that exhibited substantially elevated DNAme upon knockout and transition to 2i/L include the phosphatase *Dusp6* (60-56%), the tyrosine kinase *Jak1* (70-65%), the epigenetic regulator *Brd4* (59-51%), and the E3 ubiquitin ligase *Cop1* (56-54%). These data suggest our screen is sufficient to identify critical components of gene regulatory networks that contribute to driving complete DNA demethylation in naïve ESC.

### *Dusp6* and *Cop1* promote global DNA hypomethylation

To further investigate the role of candidates *Dusp6* and *Cop1* in epigenetic transitions, we generated independent clonal knockout ESC lines. DUSP6 is a phosphatase that acts downstream of MEK to attenuate the ERK signal cascade, whilst COP1 mediates ubiquitination and proteasomal degradation of target proteins^29,30^. We used enzymatic methyl(EM)-seq, an enhanced equivalent of bisulfite (BS)-seq, to chart the global DNA methylome in WT, *Dusp6^−/−^* and *Cop1^−/−^* naïve ESC, which confirmed mutant lines remain hypermethylated in in 2i/L (*Dusp6^−/−^* 67%; *Cop1^−/−^* 58%) (Fig 2D). Notably, elevated DNAme is distributed equivalently across genomic features such as promoters, repeats and intergenic regions, indicating a general impairment to DNA demethylation rather than failure in locus-specific resetting (Fig 2D-E).

Mechanistically, both *Cop1^−/−^* and *Dusp6^−/−^* ESC exhibited transcriptional upregulation of the *de novo* methylation machinery (*Dnmt3a*, *Dnmt3b*, *Dnmt3L*), whilst *Dusp6^−/−^* cells additionally downregulate *Stella*, which together may contribute to impaired global DNA demethylation (Fig 2F). Consistent with elevated DNAme in mutants, we observe strong repression of DNA methylation-dependent (germline) genes, whilst naïve genes are largely unaffected implying no underlying change to pluripotency networks (Fig 2F). However, we did observe inappropriate expression of some early developmental genes in *Cop1^−/−^* and *Dusp6^−/−^* ESC, and their transcriptomes clustered separately by PCA, which may partly reflect their disrupted epigenetic state (Fig 2G). In the case of *Dusp6*, impaired DNA demethylation is potentially linked with the increased ERK activity, similar to *Prdm14* mutants^31^. Taken together, our screen identifies and validates genes and pathways involved in promoting genome-scale DNA methylation transitions.

### *Dppa2/4* protect against aberrant *de novo* DNA methylation

Our screen is designed to identify both global and focal epigenome regulators. Amongst the *e*RGM screen hits, the paralogues *Dppa2* and *Dppa4* (hereafter *Dppa2/4*) consistently ranked in the top five, suggesting a role in modulating epigenome dynamics. However, in contrast to other candidates (e.g. *Dusp6*, *Cop1*, *Jak1*), deletion of *Dppa2* or *Dppa4* did not affect genome-scale DNA demethylation (Fig 2C). This could imply that, rather than a global influence, *Dppa2/4* modulate the methylation landscape at specific genomic features in pluripotent cells. Notably, the embryonic expression pattern of *Dppa2/4* coincides with pluripotency phases and consistently, they are active only in naïve (ESC) and formative (EpiLC) pluripotent cells *ex vivo*, but not in differentiated cells (Fig 3A). Nonetheless, forced expression of *Dppa2/4* in somatic cells enhances iPS cell generation through chromatin decompaction, whilst knockout mice exhibit defects in tissues that do not express *Dppa2/4*, implying an epigenetic function^32,33^.

**Figure 3.**
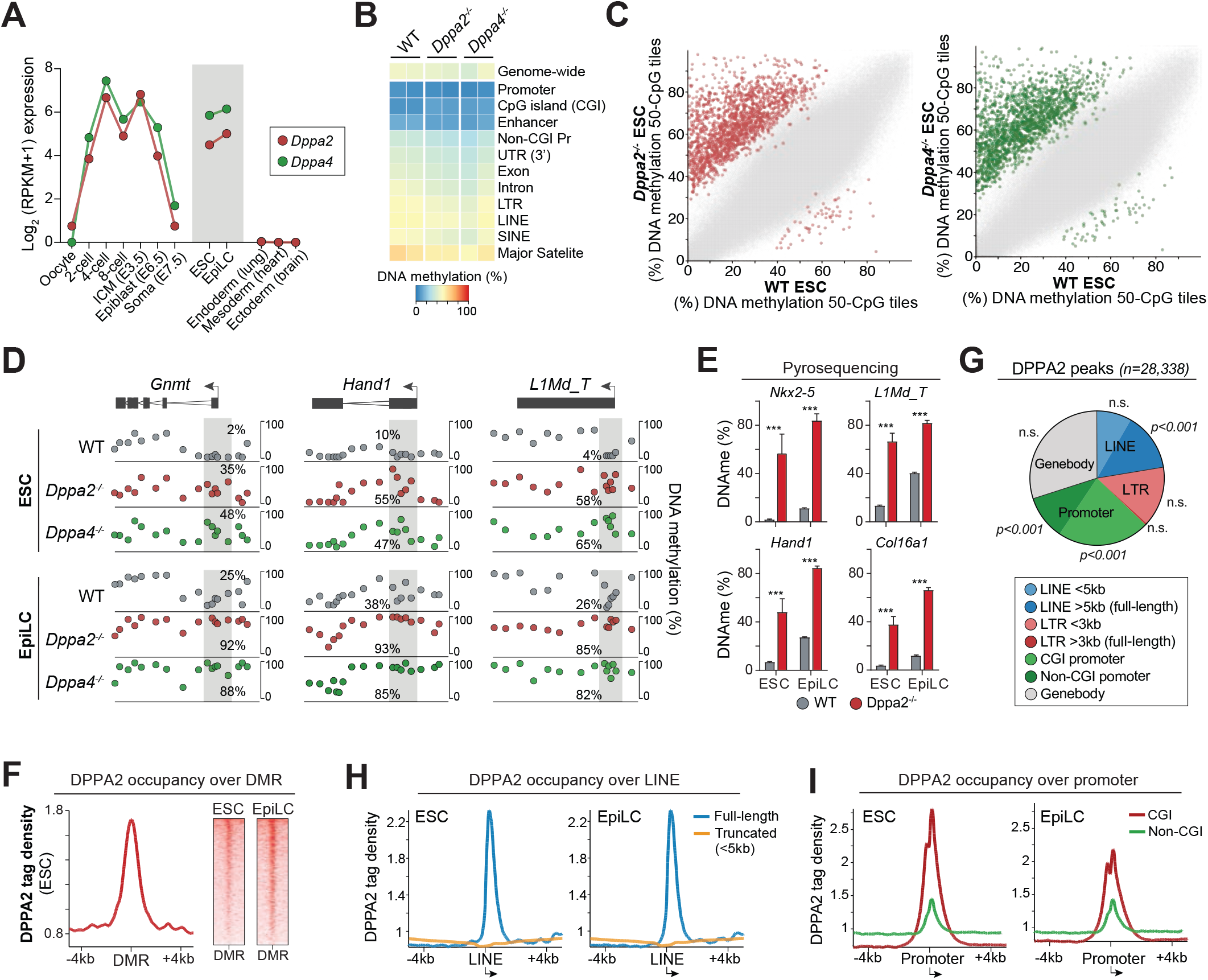
*Dppa2* and *Dppa4* occupancy promotes DNA hypomethylation in pluripotent cells. **A.)** Line plot of *Dppa2* and *Dppa4* expression dynamics during early development, in pluripotent ESC and EpiLC, and in representative somatic tissues. **B.)** Heatmap showing DNA methylation levels genome-wide and at indicated genomic features in WT, *Dppa2-/-* and *Dppa4-/-* naïve ESC. **C.)** Scatter plot of genome-wide DNA methylation at sliding 50-CpG tiles. Significant differentially methylated regions (DMR) are shown in red (*Dppa2*) or green (*Dppa4*). **D.)** Genome views showing DNA methylation distribution at genes and at LINE1 in WT, *Dppa2-/-*, *Dppa4-/-* ESC and EpiLC. Each datapoint corresponds to average methylation at sliding tiles of 15 or 20 CpGs. **E.)** Bisufite pyrosequencing quantification of DNA methylation at selected gene and *L1Md_T* promoters. Error bars are standard deviation of two biological replicates, each of multiple CpG sites. **F.)** Distribution of DPPA2 occupancy (CUT&RUN-seq) over differentially methylated regions (DMRs) +/− 4kb **G.)** Pie plot showing the genomic features associated with DPPA2 binding. **H-I.)** DPPA2 binding at **H.)** CpG-island (CGI) and non-CGI promoters and **I.)** full-length or truncated LINE1 promoters +/− 4kb. Student t test, adjusted p-value (Holm-Sidak), ns *p>0.05, * p<0.05, ** p<0.01, *** p<0.001*.

To explore this function of *Dppa2/4* molecularly we generated multiple *Dppa2^−/−^* or *Dppa4^−/−^* ESC lines, and used EM-seq to chart their global DNA methylome. Consistent with LUMA, *Dppa2/4* knockouts did not impact the global DNA demethylated status of naïve ESC (Fig 3B). However, using sliding 50-CpG genomic tiles we identified 1662 differentially methylated regions (DMR) (*logistic regression adjusted p<0.05 & binomial test p<0.01*). Remarkably 1605 of these DMRs (>96%) correspond to loci with elevated DNA methylation, whilst in contrast only 57 DMRs (3.4%) exhibit reduced DNAme, suggesting a general acquisition of focal hypermethylation in *Dppa2^−/−^* ESC (Fig 3C). *Dppa4^−/−^* ESC exhibit a highly correlated pattern of hypermethylated DMRs as *Dppa2^−/−^* (Fig 3C), likely because disruption of one protein led to reciprocal destabilization of the other (Fig S3A). Notably, DNA hypermethylation was apparent at many gene promoters that usually remain strictly unmethylated at all developmental stages (Fig 3D). This indicates that rather than impaired DNA demethylation *per se* in *Dppa2/4* mutants, there is aberrant *de novo* methylation activity that could establish DNA methylation ‘epimutations’.

To investigate this further, and determine whether such epimutations persist during differentiation, we profiled epiblast-like cells (EpiLC), which correspond to a formative state that has undergone epigenetic programming (genomic re-methylation). We observed the hypermethylated sites established in *Dppa2^−/−^* or *Dppa4^−/−^* ESC are retained in EpiLC, whilst a striking number of additional loci also acquire aberrant *de novo* methylation (DMR) in EpiLC, including promoters (Fig S3B). Indeed, direct analysis identified 354 differentially methylated promoters (DMP) in *Dppa2/4* mutants, most of which correspond to gain of DNAme. Gene ontology analysis revealed that DMPs are specifically enriched for developmental processes (*multicellular organism development FDR=0.0053*; *developmental process FDR=0.024*, *anatomical structure development FDR=0.01*) (Fig S3C). For example, *Hand1*, *Tnxb*, and *Gnmt1* are key nodes for lineage-restricted cells, and usually remain demethylated in all tissues, but acquire significant promoter hypermethylation (>90%) in *Dppa2^−/−^* and *Dppa4^−/−^* ESC and EpiLC (Fig 3D & S3D). Intriguingly, in addition to developmental gene promoters we observed that DMR from *Dppa2/4*-mutants are enriched specifically for the 5’ end of *full-length* (>5kb) LINE1 elements (indicative of evolutionarily young LINEs), but not for truncated LINEs and LTR elements (Fig S3B). Indeed, direct analysis identified 1131 differentially methylated LINE1 (DML), of which over 80% are *L1Md_T* (Fig 3D). We used pyrosequencing to verify that LINE (*L1Md_T*), as well as the promoters of *Hand1*, *Nkx2-5* and *Col16a1*, acquire DNA methylation in *Dppa2^−/−^* naïve ESC (Fig 3E), confirming a disrupted epigenomic landscape.

Because the DMRs are focal rather than global, we next asked whether they reflect localised DPPA2/4 activity. To test this, we performed CUT&RUN for DPPA2 binding in WT cells and observed genomic occupancy is strikingly increased over sites that become hypermethylated DMR in *Dppa2/4^−/−^* cells, suggesting DPPA2 may act proximally to sculpt the DNA methylome (Fig 3F). Indeed, DPPA2 binding peaks (n=28,338) are significantly enriched specifically over gene promoters (*p*<0.001) and the 5’ end of full-length LINE1 elements (>5kb) (*p*<0.001) (Fig 3G), consistent with DMR associations. Overall, promoters and LINE account for >65% of DPPA2 genomic occupancy. The latter enrichment is specific for full-length LINE since DPPA2 is not enriched at truncated LINEs or other repetitive genomic features (Fig 3H). Amongst promoters, DPPA2 exhibits a preference for CpG-dense loci, which is reflected by its GC-rich binding motifs and preference for CpG island (CGI) promoters (Fig 3I & S3E). Notably, DPPA2 binding was observed at the sensor region used for *e*RGM (Fig S3F), explaining why a focal DNAme modulator was identified in the screen. In summary, *Dppa2* and *Dppa4* have a non-redundant role in protecting a target subset of developmentally-associated promoters and full-length LINE elements from acquiring *de novo* DNA hypermethylation during naïve and formative pluripotency phases, when *Dppa2/4* are specifically expressed.

### DPPA2/4 binding establishes a permissive chromatin state

To understand the broader chromatin features associated with DPPA2 occupancy, and susceptibility to hypermethylation, we used CUT&RUN to profile H3K4me3, H3K27me3, and H3K9me9. Notably, we observed that DPPA2 occupancy directly correlates with strong H3K4me3 enrichment in WT cells, across all binding sites (Fig S4A). H3K27me3 is also enriched at a subset of DPPA2-bound sites, establishing bivalent states, but H3K9me3 is largely depleted. Strikingly, the H3K4me3 enrichment at DPPA2-bound promoters occurs irrespective of expression state, in both ESC and EpiLC (Fig 4A), implying that DPPA2 may directly target H3K4me3, rather than H3K4me3 reflecting expression status of DPPA2-bound sites. Importantly, loci that acquire hypermethylation in *Dppa2/4* mutants (DMR) correspond to genomic regions that are H3K4me3-enriched and DPPA2-bound in WT (Fig S4B). Taken together this suggests a potential connection between DPPA2 occupancy, H3K4me3 and DNA methylation status.

**Figure 4.**
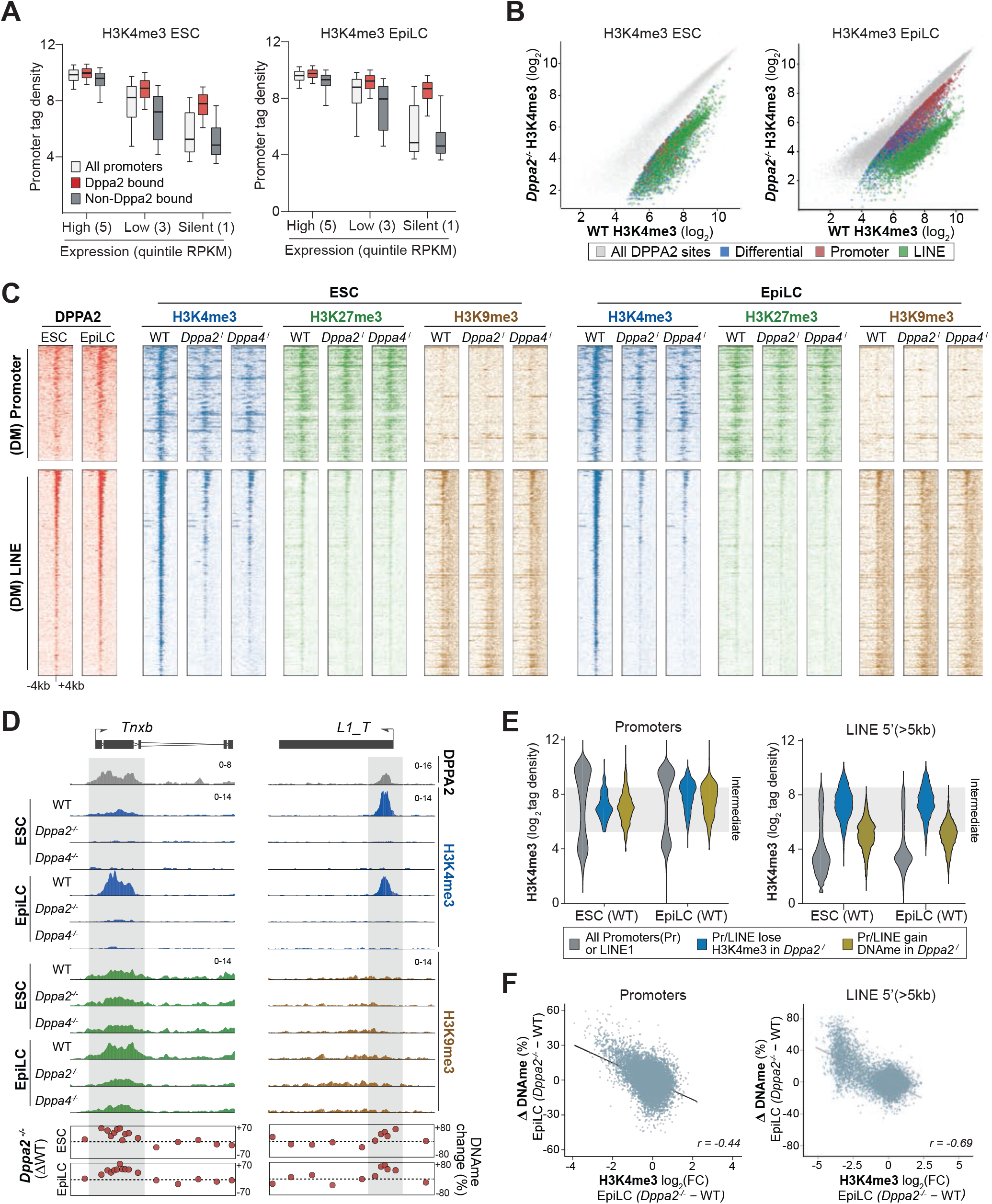
*Dppa2* and *Dppa4* establish chromatin states to safeguard against *de novo* methylation. **A.)** Boxplot showing H3K4me3 enrichment over all promoters, DPPA2-bound, and non-DPPA2-bound promoters, binned for expression quintile (RPKM). **B.)** Scatter plot of H3K4me3 enrichment at DPPA2-binding sites in WT and *Dppa2−/−* ESC and EpiLC. **C.)** Density plot revealing enrichment of H3K4me3, H3K27me, and H3K9me3 (Cut&RUN-seq) centered on differentially methylated promoters (DMP) (upper) or LINEs DML) (lower) +/− 4kb. Plots are ordered by DPPA2 binding enrichment (left). **D.)** Representative genome view of a developmental promoter and a LINE1 with tracks for DPPA2 occupancy, H3K4me3, H3K27me3 or H3K9me3, and changes in DNA methylation in WT, *Dppa2−/−* and *Dppa4−/−* cells. **E.)** Violin plot detailing distribution of H3K4me3 in WT cells at all promoters/LINE1 and those susceptible to H3K4me3-loss or DNAme gain upon *Dppa2* knockout **F.)** Scatter plot showing correlated inter-relationship between changes in H3K4me3 and DNA methylation upon *Dppa2* knockout at all genes (left) and all full-length LINE1 (right).

To investigate this further we assayed H3K4me3, H3K27me3 and H3K9me3 in *Dppa2^−/−^* and *Dppa4^−/−^* ESC and EpiLC. Remarkably, deletion of *Dppa2* or *Dppa4* results in a dramatic loss of H3K4me3 across a significant subset of DPPA2-bound sites in both ESC and EpiLC, whilst the remaining sites are apparently unaffected (Fig 4B). Notably, the subset of DPPA2 sites that lose H3K4me3 are enriched for full-length LINE1 elements and promoters (Fig 4B). Moreover, the effect on H3K4me3 is specific, since there is no significant change of H3K9me3 in mutants, whilst H3K27me3 is reduced at some loci such as *Txnb*, but relatively unaffected at most (Fig 4C-D & S4D). In general, the loci that lose H3K4me3 and gain DNAme in *Dppa2/4*-mutants are associated with specific absolute levels of H3K4me3 enrichment in WT cells; intermediate for promoters and (relatively) high for full-length LINE1 (Fig 4E). Because H3K4me3 can directly impair *de novo* DNA methylation via inhibiting *Dnmt3L* access ^34^, the dramatic depletion of H3K4me3 in *Dppa2/4*-mutants may enable aberrant DNA hypermethylation. In support of this, the differentially methylated promoters (DMP) and LINE1 (DML) correspond to the loci that exhibit reduced H3K4me3 (Fig 4C-D). Furthermore, by investigating all promoters and LINE1, we observe a striking negative correlation (*p*<2.2e-16) between the level of H3K4me3 loss and of DNAme gain upon *Dppa2* KO (Fig 4F).

In summary, abrogation of *Dppa2/4* is linked with depletion in H3K4me3 at a specific subset of DPPA2-target loci, which directly correlates with acquisition of aberrant DNA hypermethylation. *Dppa2/4* could therefore integrate chromatin states to safeguard the pluripotent epigenome, particularly at developmental-associated genes and LINE1 elements.

### *Dppa2/4* ensure a competent epigenome for developmental expression

To understand the relevance of *Dppa2/4*-mediated epigenome surveillance for developmental competence, we assessed the transcriptome. *Dppa2/4^−/−^* ESC express an unperturbed naive pluripotency network, and undergo apparently normal exit from pluripotency, since naïve markers (*Nanog*, *Klf2*, *Prdm14* etc) are downregulated equivalently to WT EpiLC whilst formative markers (*Fgf5*, *Wnt3*, *Dnmt3a*) are appropriately upregulated (Fig 5A). Moreover, there is no difference in expression of the DNA methylation machinery or chromatin-modifying genes, which taken together implies *Dppa2/4* do not have an overarching influence on naive, early differentiation or epigenome gene regulatory networks (Fig 5A).

**Figure 5.**
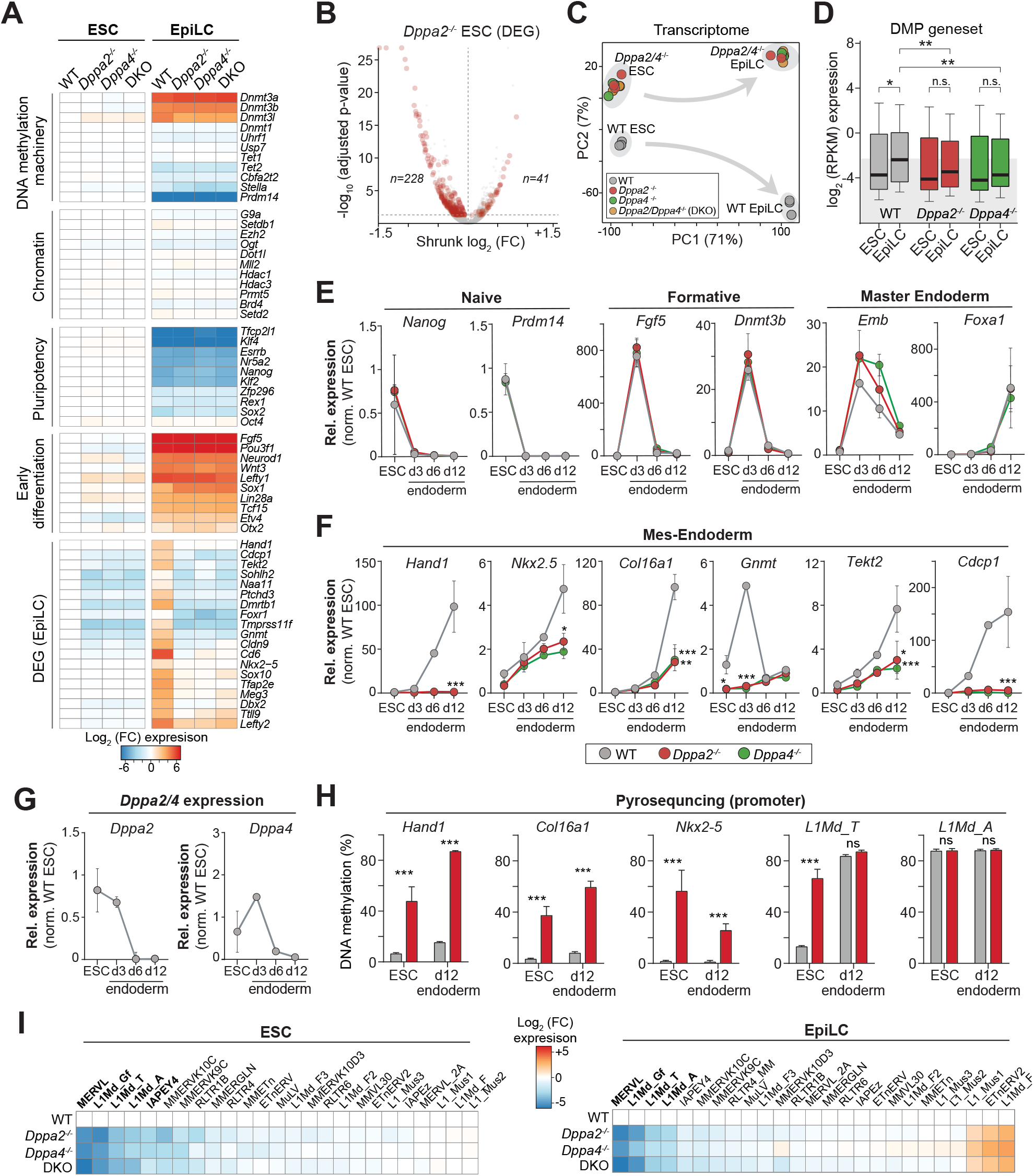
Developmental genes and LINE1 acquire transcriptional silencing memory in *Dppa2/4*-mutants. **A.)** Heatmap showing expression changes (RNAseq) for selected pathways in WT, *Dppa2−/−, Dppa4−/−* and DKO. **B.)** Volcano plot highlighting significant differentially expressed genes (DEG) from *Dppa2−/−* ESC. **C.)** Principal component analysis of global transcriptomes. **D.)** Boxplot showing activity of genes linked with a differentially methylated promoter (DMPs) in WT, *Dppa2−/−* and *Dppa4−/−* cells. Box indicates the 25^th^, median and 75^th^ percentiles, whiskers the 10^th^ to the 90^th^ percentiles. Significance by students t-test, ns p>0.05, * p<0.05, ** p<0.01. **E-G.)** qRT-PCR quantification of expression of selected genes during ESC to endoderm differentiation. **H.)** Bisulfite pyrosequencing quantification of DNA methylation at selected promoters and LINE1. Error bars are standard deviation of two biological replicates, each of multiple CpG sites. **I.)** Heatmap showing expression of all full length LINE1 (>5kb) and LTR (>3kb) elements above threshold in WT, *Dppa2−/−* and *Dppa4−/−* ESC and EpiLC. Significance by adjusted students t-test (Holm-Sidak), ns p>0.05, * p<0.05, ** p<0.01, *** p<0.001.

Globally, *Dppa2/4^−/−^* naïve ESC exhibit a distinct but broadly comparable gene expression signature to WT, with 269 and 245 differentially expressed genes (DEG) in *Dppa2* and *Dppa4* knockout respectively, the majority of which (85% and 82%) are downregulated (Fig 5B & S5A). Induction of EpiLC leads to a more divergent transcriptome as judged by principal component analysis (Fig 5C), with 801 and 611 differential genes, again preferentially downregulated. Significantly, gene ontology indicated these DEG in EpiLC specifically relate to developmental processes (*single-multicellular organism process* FDR *=0.000004*, *cell differentiation* FDR *=0.00012*) (Fig S5B), which reflects a general failure to activate genes involved in lineage-specific functions, particularly mesendoderm regulators. For example, *Hand 1*, *Cldn9*, *Tnxb* and others all fail to initiate primed expression in mutant EpiLC (Fig 5A & S5C). This could be linked with the ectopic promoter DNA methylation acquired in the preceeding ESC state. Indeed, the DMP geneset collectively (*n=354*), which comprises many of the same mesendoderm genes including *Hand1*, *Tnxb*, *Ttl9*, *Cldn9* and *Gnmt*, is significantly upregulated in WT EpiLC (*p*=0.018) consistent with priming developmental genes, but strikingly, fails to initiate activation in either *Dppa2^−/−^* (*p*=0.29) or *Dppa4^−/−^* EpiLC (*p*=0.40), suggesting they have lost competence for expression (Fig 5D).

To understand if the impaired expression of mesendoderm genes in EpiLC represents a delay in their activation or stable silencing, we induced endoderm differentiation for 12 days. *Dppa2^−/−^* cells appeared morphologically equivalent to WT and activated master endoderm regulators such as *Emb* and *Foxa1* with comparable dynamics, indicating no general impairment in differentiation (Fig 5E). However, endoderm-associated genes including *Gnmt*, *Nkx2-5*, *Col16a1* and *Hand1* all exhibited a highly significant failure to activate in mutants, even after 12 days of endoderm induction, implying an absolute blockade in their response (Fig 5F). Importantly, *Dppa2* is rapidly downregulated after 3 days of endoderm differentiation, but impaired gene upregulation manifests at later timepoints, suggesting a memory of DPPA2 prior activity (Fig 5G). Indeed, pyrosequencing revealed ectopic promoter DNA methylation established in ESC propagates through to d12 endoderm (Fig 5H). Together this indicates that the absence of *Dppa2/4* in pluripotent phases leads to impaired competence for gene activation during later differentiation. Importantly, cell fate transition *per se* appears unperturbed, but rather specific genes within the developmental programme are rendered stably epigenetically silenced.

We next asked whether repetitive element activation is also affected by *Dppa2/4*, since many evolutionary young LINE1 become hypermethylated and lose H3K4me3 in their absence (Fig 3D & 4C). We observed a particularly striking downregulation of full-length (>5kb) LINE families (*L1Md_T*, *L1Md_A* and *L1Md_Gf*) in *Dppa2/4*-mutant ESC and EpiLC (Fig 5I & S5D). Indeed, we confirmed this with independent qRT-PCR showing that disruption of *Dppa2* in ESC and EpiLC leads to extensive repression of *L1Md_T* (Fig S5E), whilst *MERVL* is also strongly repressed, consistent with recent reports^35^. *IAP* and other LTR elements were largely unaffected. These data suggest that the same *Dppa2/4*-depedent system that maintains epigenetic competence at developmental promoters may have been co-opted by LINE1 elements to evade epigenetic silencing in pluripotent phases.

### H3K4me3 and DNAme interact to confer functional epigenetic memory

We finally investigated whether induced DNA methylation and H3K4me3 ESC loss is functionally instructive for the subsequent gene silencing memory. We noted that depletion of promoter H3K4me3 and gain of DNAme are both correlated with gene repression in *Dppa2^−/−^* cells (Fig S6A). Moreover, altered DNAme and H3K4me3 are also directly anti-correlated (Fig 4F), implying a hierarchy of robust molecular changes upon *Dppa2/4* abrogation. To determine if the acquired DNA methylation (and H3K4me3 loss) could instruct gene repression, we deleted *Dnmt1* in *Dppa2^−/−^* ESC, to generate compound mutants (*Dnmt1^−/−^*, *Dppa2^−/−^*) that are hypomethylated and predicted to erase the ectopic DNAme at developmental promoters and LINE1. Remarkably, analysis of the DMP geneset that acquire aberrant promoter methylation and silencing in *Dppa2^−/−^* EpiLC, revealed that additional deletion of *Dnmt1* partially rescues their activation block in EpiLC. This effect is significant (*p*=0.024) among genes with CpG island (CGI) promoters, but not non-CGI promoters (*p*=0.23) (Fig 6A). Moreover, we observed re-activation of *L1Md_T* elements in compound *Dnmt1^−/−^, Dppa2^−/−^* ESC and EpiLC (Fig 6B). These data imply that ectopic DNAme in *Dppa2^−/−^* cells is instructive, at least at some CpG-dense genes and LINE1, and directly impairs their response to inductive activating signals.

**Figure 6.**
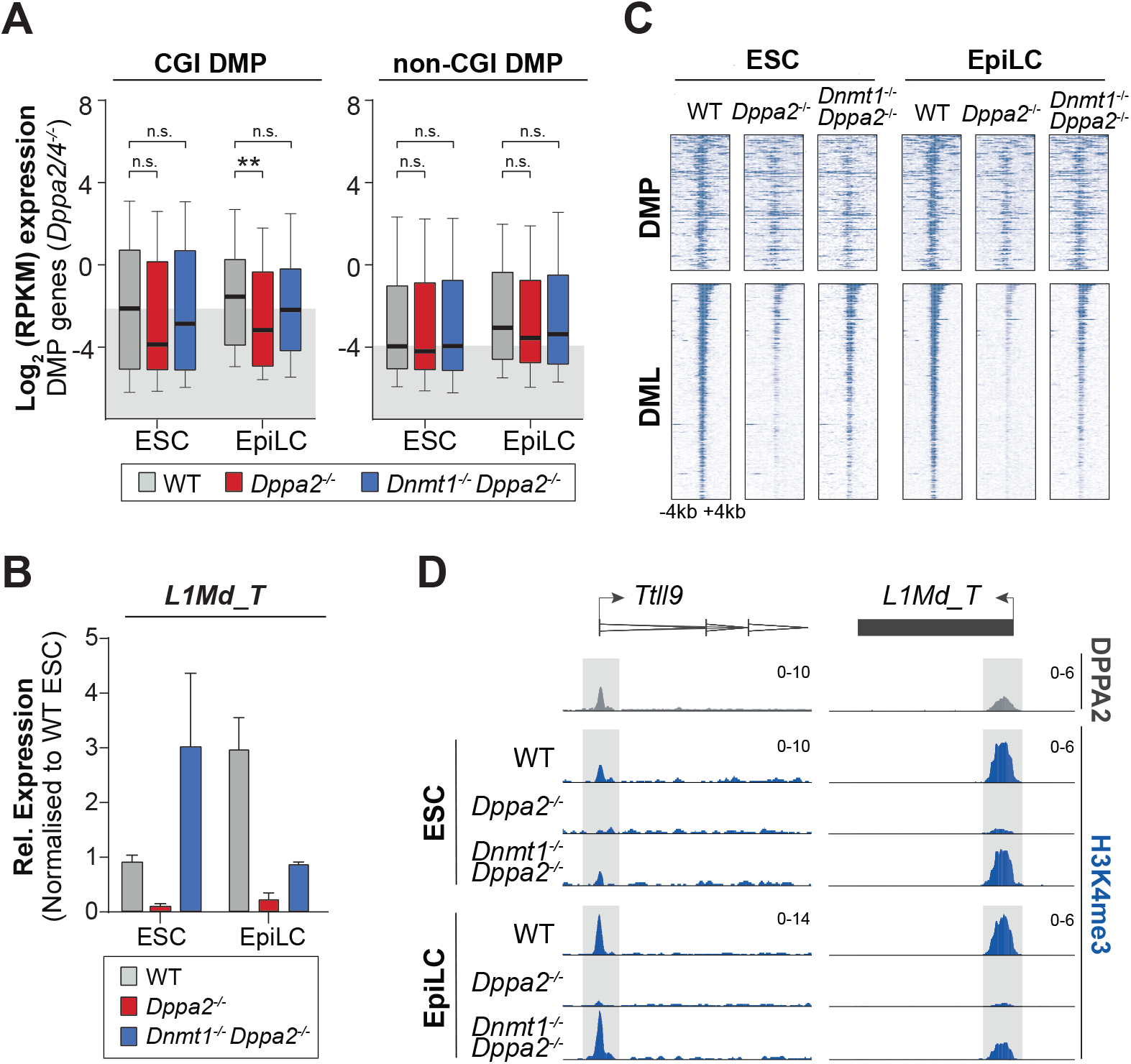
Functional interaction between DNAme, H3K4me3, and gene silencing. **A.)** Boxplot showing expression of all genes associated with differentially methylated promoters (DMPs) in WT, *Dppa2−/−* and rescue compound-*Dnmt1−/−, Dppa2−/−* cells. Box indicates the 25^th^, median and 75^th^ percentiles, whiskers the 10^th^ to the 90^th^ percentiles. Significance by students t-test, ns p>0.05, * p<0.05, ** p<0.01. **B.)** qRT-PCR quantifying expression of *L1Md_T* in WT, *Dppa2−/−* and compound-*Dnmt1−/−, Dppa2−/−* cells, normalised to WT ESC. **C.)** Density plot showing enrichment of H3K4me3 centered on differentially methylated promoters (DMP) (upper) or LINEs (DML) (lower) +/− 4kb. **D.)** Representative H3K4me3 genome tracks of a developmental promoter and LINE1 (*L1Md_T*) instance.

We next investigated whether the depletion of H3K4me3 in *Dppa2-*mutant cells is also affected in *Dnmt1-Dppa2*-compound mutants and surprisingly observed a reinstatement of H3K4me3 at a subset of promoters and most LINE1 elements (Fig 6C-D). This indicates a potentially complex interplay whereby absence of *Dppa2/4* leads to loss of H3K4me3, enabling aberrant DNA methylation, but that subsequently removing ectopic DNAme tips the balance back, allowing H3K4me3 to re-accumulate through *Dppa2/4*-independent mechanisms (Fig S6C). More generally, we show altered DNAme and H3K4me3 in the absence of *Dppa2/4* can propagate through differentiation to manifest as instructive gene silencing at future developmental stages, long after the ‘epimutation’ is established. *Dppa2/4* therefore act as a safeguarding system during dynamic epigenome remodelling phases to ensure epigenetic competence for impending multi-lineage development.

## DISCUSSION

Here, we have established a ratiometric reporter of DNA methylation (*e*RGM) that is enhanced to enable unbiased CRISPR screening. By coupling this with an ESC model of developmental DNAme reprogramming, we identify and validate epigenome modulators that influence both global DNA demethylation events and also focal DNAme states (*Dppa2* & *Dppa4*). The global regulators relate to diverse pathways such as m6A RNA methylation (e.g. *Mettl3*, *Virma* and *Ytdhf2*), LIF signaling (e.g. *Lifr*, *Jak1*, *Stat3*) and E3 ubiquitin ligases, which presumably exert influence through acting as upstream regulators, as recently reported for *Nudt21*^36^. Amongst these, we show the phosphatase *Dusp6* is necessary for completion of global DNA demethylation in naïve ESC. DUSP6 functions to attenuate MEK/ERK signaling^29^, which is linked with DNA methylation^22,37^, suggesting a likely connection. Indeed X-linked DUSP9 contributes to female-specific ESC hypomethylation by influencing MEK/ERK ^38^, and DUSP6 could play a comparable but non-redundant role in thresholding MEK/ERK more generally in pluripotent cells to promote epigenome erasure. Interestingly another screen hit, *Med24* (rank=3), was also found to impact MEK/ERK signaling^39^. Mechanistically, *Dusp6* may function via modulation of the *de novo* methylation machinery, since *Dnmt3a/b/l* are known MEK/ERK targets and consistently, are upregulated in *Dusp6^−/−^* cells here. The mechanism through which another hit, the E3 ubiquitin ligase *Cop1*, modulates global epigenetic state is less clear, but could relate to regulating the stability of proteins involved in maintaining DNAme, such as UHRF1^10^.

In addition to global regulators we identify *Dppa2/4*, which we show guard against ectopic *de novo* methylation activity at key genomic sites during phases of both DNAme erasure (in naïve ESC) and re-methylation (EpiLC). This *Dppa2/4* activity represents an important complementary pathway to genome-scale (re)programming mechanisms, because it ensures a permissive epigenetic state at key lineage-associated genes, rendering them competent to respond to future activating signals. Previous studies have shown that *Dppa2/4* overexpression enhances iPS cell generation, and they are linked with facilitating the 2-cell programme in ESC via modulating *Dux*, suggesting broad functional roles^32,35,40^. Nevertheless, *Dppa2/4*-mutant mice undergo normal embryogenesis but die perinataly due to aberrant gene repression in lung, where *Dppa2/4* are not expressed^33^, implying the phenotypically-relevant activity of *Dppa2/4* is ensuring lineage-associated genes are appropriately primed during early development. We dissect this molecularly by demonstrating absence of *Dppa2/4* leads to a striking loss of H3K4me3 and parallel acquisition of *de novo* DNA methylation at key developmental genes and LINE1 elements, which propagates to manifest as epigenetic silencing in lineage-restricted cells. The marked equivalence of DPPA2 or DPPA4 in this function likely reflects that they reciprocally stabilize each other (Fig S3), whilst the unusual association of DPPA2 outside of classical pluripotency networks could underpin lack of redundancy more generally^41^.

Mechanistically, several lines of evidence suggest that DPPA2 targets H3K4me3. First DPPA2 genomic binding sites are highly H3K4me3-enriched, irrespective of underlying transcription. Second H3K4me3 is lost at a subset of sites upon *Dppa2*-deletion, and third, DPPA2 is reported to interact with the H3K4me methylase MLL2^42^. Because H3K4me3 restricts the recruitment of *de novo* DNA methyltransferases^34,43^, the depletion of H3K4me3 at these loci may enable access for ectopic DNA methylation to follow^44^. Indeed, we observe a striking correlation between degree of H3K4me3 loss and DNAme gain in *Dppa2/4* mutants. Consistently, a *Dnmt3a* engineered to tolerate H3K4me3 enables aberrant *de novo* methylation at developmental genes^45^, supporting a model whereby *Dppa2*-dependent H3K4me3 protects against DNAme. Such a system is likely necessary due to widespread *de novo* methylation activity throughout developmental (re)programming phases^15^, which presumably maintains epigenetic repression at specific genomic compartments such as TE, but could also aberrantly target developmental genes if not restrained^46^. Notably, by counteracting *de novo* methylation DPPA2/4 may also facilitate H3K27me3 accumulation and bivalency, because a subset of loci exhibit H3K27me3 depletion in *Dppa2/4*-mutants. This could reflect direct loss of targeting by DPPA2/4 or inhibition of PRC2 binding by acquired DNAme^47–49^. Functionally, the acquisition of ectopic DNAme and loss of H3K4me3 appears to be instructive for epigenetic silencing upon differentiation, at least at some CpG-dense promoters. Indeed, erasing acquired promoter DNAme partially rescues expression defects, and also reinstates H3K4me3. This suggests a switch-like interdependency whereby DPPA2-dependent H3K4me3 impairs DNAme, but acquired DNAme reciprocally prevents H3K4me3 accumulation, potentially underpinning stable transcriptional memory at target genes.

In addition to maintaining epigenetic competence at developmental genes, we observe that evolutionary young LINE1 elements (*e.g. L1Md_T*) directly rely on *Dppa2* for H3K4me3 and their activity. This may reflect a strategy of successful LINE1 elements that have acquired/co-opted DPPA2 binding sites at their 5’ ends to protect against host-directed epigenetic silencing. This would in turn enable expression of full-length LINE1 during early development when *Dppa2/4* are expressed, which aligns well with the optimal period for retrotransposition^50^. This scenario would represent a genomic conflict, whereby *Dppa2/4* activity is critical for epigenetic competence of lineage-associated genes, and therefore essential for viability, but also renders full-length LINE1s transcriptionally competent; a potential threat to genome integrity. An alternative scenario is that *Dppa2/4*-mediated activation of LINE1 reflects an important developmental role for the host. For example, LINE1 activation has been linked with establishing zygotic chromatin accessibility and also directly represses the 2-cell (2C) programme via binding Nucleolin/TRIM28^16,51^. Consequently, *Dppa2/4*-dependent LINE1 activation could represent exaptation by host systems to exploit LINE1 functionality at critical developmental points. In any case, *Dppa2/4* are placed at the centre of an epigenetic competence circuit in pluripotent cells that facilitates expression of both LINE1 and developmental genes.

In summary, we characterise upstream gene networks that influence global DNA methylation erasure, and additionally uncover a complementary pathway that protects against the counterforce of uncontrolled *de novo* methylation during (re)programming, to ensure developmental competence.

## ACKNOWLDEGMENTS

We thank Mathieu Boulard and Phil Avner for critical reading of the manuscript, and Erna Magnusdottir and Alexander Aulehla for their contributions to TAC discussions. We are grateful to Haruhiko Koseki and Jafar Sharif for kindly sharing floxed *Dnmt1* ESC. We also thank Cristina Policarpi for CUT&RUN advice, and all members of the Hackett lab for experimental support. We are grateful to all EMBL core facilities, and in particular the genetic and viral engineering (Jim Sawitzke) and flow cytometry (Cora Chaddick) facilities, for key experimental assistance. This study was funded by a European Molecular Biology Laboratory (EMBL) programme grant to J.A.H.

## AUTHOUR CONTRIBUTIONS

K.H.G performed experiments and bioinformatics analysis, and contributed to the manuscript. J.A.H designed and supervised the study, performed bioinformatics analyses and wrote the manuscript.

## COMPETING INTERSTS

We declare no financial or non-financial competing interests.

## METHODS

### Cell culture and differentiation

Murine embryonic stem cell (ESC) lines (mixed 129:B6, XY) were routinely maintained and manipulated on gelatin in <u>t</u>itrated 2i/L (*<u>t</u>*2i/L) culture media (NDIFF 227 supplemented with PD0325901 (200nM), CHIR99021 (3μM), LIF (1000 U/ml), FBS (1%), and penicillin/streptomycin; filtered through 0.22μM filter) in a humidified CO_2_ incubator at 37°C. *t*2i/L maintains genomic stability 37 and DNA hypermethylation ^22^ (Fig 1). ESC were passaged every two days or at sub-confluence by dissociation with TrpLE. To induce DNA hypomethylation ESC were transitioned into full 2i/L culture media ^52^ (NDIFF 227 supplemented with PD0325901 (1μM), CHIR99021 (3μM), LIF (1000 U/ml), FBS (1%), and penicillin/streptomycin; filtered through 0.22μm) on gelatin for up to 14 days. To induce epiblast-like cells (EpiLC), 3×10^4^ naïve ESC per cm^2^ were seeded on fibronectin coated wells in EpiLC media (NDIFF 227 supplemented with knockout serum replacement (1%) (KSR), Activin-A (20 ng/ml), bFGF (12.5 ng/ml) and penicillin/streptomycin) for 44 hours. To induce endodermal differentiation 6×10^3^ naïve ESC per cm^2^ were seeded on gelatin coated wells in 2i/L media. After overnight culture cells were washed 3x with PBS and Endoderm media ^53^ was introduced (RPMI supplemented with L-Glutamine (2mM), FBS (0.2%), IDE1 (5μM) and penicillin/streptomycin). Endodermal media was replaced every two days.

### Enhanced ratiometric RGM (*e*RGM) constructs

A H2B-GFP-SV40pA cassette (from addgene #11680) was cloned downstream of the mouse core imprinted *Kcnq1ot1* promoter (mm10, Chr7:143,296,371-143,296,745) into a piggyBac backbone vector using InFusion assembly (pPB-Kcnq1ot1-H2B-GFP). A genomic ‘DNAme sensor’ region derived from the mouse *Dazl* locus (mm10, chr17:50,293,285-50,294,435) was amplified and inserted in the *antisense* orientation upstream of the *Kcnq1ot1* promoter using infusion assembly to generate pPB-as *Dazl*sensor-*Kcnq1ot1*-H2B::GFP, which exhibits methylation-sensitive activity. To establish a ratiometric system an additional construct was generated by cloning the methylation-insensitive *EF1a* promoter upstream of an mCherry::H2B cassette into a piggyBac vector to generate pPB-EF1a-H2B::mCherry). Correct assembly and sequences were confirmed by tiled sanger sequencing and vectors were amplified and purified by endotoxin-free midi-preparations.

### Generation of *e*RGM ESC lines

ESC lines carrying floxed *Dnmt1* alleles ^23^ and WT ESC were transfected with pPB-as *Dazl*sensor-*Kcnq1ot1*-H2B::GFP (*in silico* DNA methylated with M.SssI), pPB-*EF1a*-H2B::mCherry, pPB-*spCas9*-Hygro ^54^, and PBase using Lipofectamine 3000. Transfected cells were selected for *spCas9* integration in titrated 2i/L using Hygromycin (250 μg/ml) for 5 days, and clonally-derived cell lines were subsequently isolated and expanded. Clonal ESC lines were tested to confirm single-copy integrations by qPCR on genomic DNA, and their response to DNA demethylation was confirmed by evaluating *e*RGM (GFP and mCherry) expression using flow cytometry after culture for 7 days in *t*2i/L (hypermethylated) or 2i/L (hypermethylated). Further confirmation of *e*RGM response was determined by addition of tamoxifen (TAM) (800nM) for 6 days in *t*2i/L to induce conditional *Dnmt1* knockout and DNA hypomethylation. The clonal ESC lines exhibiting the best dynamic range of *e*RGM response were selected and two independent lines (*e*RGM #1 and *e*RGM #2) were used for CRISPR screening.

### CRISPR Screen

**Lentiviral particles carrying the Brie gRNA library** ^24^ were produced by transfecting Lenti-X HEK 293T with *pPax2* plasmid, *pMD2.G* plasmid, and the *Brie* library plasmid with lipofectamine 3000 in a BSL2 tissue-culture facility. Lentivirus-containing supernatant (media) was harvested at 48 and 72 hours after transfection, and clarified by filter through a 0.22μm low protein-binding unit. Viral particles were concentrated using Lenti-X concentrator, according to manufactures instructions, and resuspended in NDIFF 227. Lentiviral activity and efficiency were determined by transducing ESC across a titration curve, and assaying cell survival following puromycin selection for viral encoded integration of a resistance cassette. To generate KO library cell lines, 7×10^7^ ESC of eRGM #1 and eRGM #2 cultured in *t*2i/L were transduced with a pre-determined number of lentiviral particles carrying the Brie genome-wide CRISPR knockout sgRNA library (*n=78,637*)^24^ to ensure ~45% infection efficiency (>400 fold gRNA coverage). Transduced cells were selected for with puromycin (1.2μg/ml) for 7 days in *t*2i/L. The minimum population of cells was maintained at > 3.2×10^7^ during passaging (>400 fold coverage) to ensure maintennace of library coverage. To initiate the screen KO library *e*RGM cell lines were transitioned into 2i/L for 12 days to drive extensive DNA hypomethylation. At 12 days GFP-negative cells (defined as the lowest 1% of GFP-expression) that also maintained normal mCherry expression - together indicative of incomplete epigenetic resetting - were purified by flow cytometry (291,248 and 237,121 for *e*RGM#1 and *e*RGM#2, respectively). We additionally collected total unsorted cells (>3×10^7^) from both *t*2i/L and 2i/L, and GFP-positive cells (top 1%) from *t*2i/L (indicative of loss of epigenetic silencing), as controls. Genomic DNA was isolated from purified populations using the Quick-DNA microprep plus kit (Zymo Research, #D3020) or a DNeasy blood and tissue kit (Qiagen, #69504). Integrated gRNAs from each population were amplified from genomic DNA using custom primers with the P7 flow cell overhangs: 5’-CAAGCAGAAGACGGCATACGAGATNNNNNNNNGTGACTGGAGTTCAGACGTGTGCTCTTCCGATCTTCTACTATTCTTTCCCCTGCACTGT-3’ (8bp Barcode) and P5 overhang: 5’-AATGATACGGCGACCACCGAGATCTACACTCTTTCCCTACACGACGCTCTTCCGATCTTTGTGGAAAGGACGAAACACCG-3’ using Q5 Hot Start High-Fidelity polymerase (NEB, #M0494S) for 21-24 cycles. sgRNA amplicons were purified with SPRI beads (Beckman Coulter, #B23318), following the manufacturer’s instructions, and dsDNA was quantified with Qubit III. Amplicon libraries were multiplexed and SE50 sequenced with a Nextseq500 Illumina system.

### Gene editing in ESC

To generate clonal knockout (KO) lines with CRISPR/Cas9, ESC carrying *e*RGM and maintained in *t*2i/L were transiently transfected with a vector carrying a gRNA cassette targeted against a critical coding exon of the gene of interest (*Dppa2*, *Dppa4*, *Dusp6*, *Cop1* etc), and selected with puromycin (1.2μg/ml) for 60 hours. Transfected cells were subsequently seeded at low density (1000 cells per 9.6cm^2^) for single clone isolation. After clonal expansion, successful homozygous knockout lines (carrying frame-shifting indels) were confirmed by Sanger sequencing using the Tracking of Indels by DEcomposition (TIDE) tool ^55^, by western blot, and via functional assays. To generate population scale KO of multiple candidate factors (*n=24*), gRNAs targeting the gene(s) of interest were cloned into a piggyBac vector containing the enhanced gRNA cassette ^56^. This was co-transfected with PBase into independent *e*RGM lines that carry *spCas9* activity using lipofectamine 3000, following the manufacturers recommendations. ESC were selected for successful integration of the gRNAs for 7 days with puromycin (1,2μg/ml), which drives iterative targeting of the gene of interest until indel formation is induced. We assayed the population for successful KO by western blot and flow cytometry and typically observed >95% of individual cells within each population carried homozygous functional knockout KO.

### Flow Cytomtery

Cells were gently dissociated into single cell suspension using TrpLE and resuspended in PBS + 1% FBS (FACS media) and filtered. Fluorescent activated cell sorting (FACS) was performed using a FACS Aria III (Becton Dickinson) and FACS Diva software. For flow analysis samples were run on Attune NxT (ThermoFisher). Data were analysed using FlowJo v10.5.3 (Tree Star, Inc.).

### LUMA

The LUminometric DNA Methylation Assay (LUMA) was used to measure global DNA (CpG) methylation levels. Briefly 200-500ng of purified genomic DNA was split equally and subjected to two parallel 4 hour restriction digests at 37°C. Digestion A: *HpaII/EcoRI*, and Digestion B: *MspI/EcoRI*, whereby *EcoR1* is included as an internal reference. An equal volume of annealing buffer was added and samples were loaded into a PyroMark Q24 Advanced pyrosequencer to quantitate the protruding ends from each digestion, using the dispensation order GTGTGT<u>CA</u>CACA<u>GT</u>GTGT. The % DNA methylation was calculated by comparing the *EcoRI* normalized *HpaII* signal intensity ratio to the normalized *MspI* signal intensity ratio using the formulas:

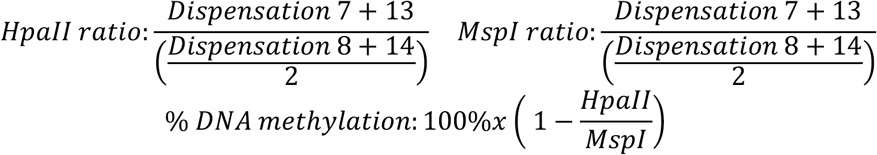

### Western Blot

Cellular protein was extracted using RIPA buffer (Sigma, #R0278) with protease inhibitors (Roche) at 4°C for 30 minutes. After centrifuging at full speed, cell lysis supernatant was collected and Bolt LDS sample buffer (Thermo Fisher, B0007) and bolt reducing agent (Thermo Fisher, B0004) were added to the samples These were heated at 70°C for 10 minutes and loaded onto 4-12% Bis-Tris gel (Thermo Fisher, NW04125BOX). The proteins were separated with 150V electrophoresis for 30 minutes and blotted onto a PVDF membrane using the iBlot Dry 2 blotting system. The membrane was blocked in 5% milk/PBS for 1 hour at RT followed by incubation with primary antibody (1:500 to 1:1000 dilution) with 5% milk/PBS at 4°C overnight with agitation. After washing twice with PBS/0.1% Tween the membrane was incubated for 1 hour at RT with a HRP-linked secondary antibody diluted 1:10,000 in 5% milk/PBS. The membrane was washed thrice with PBS 0.1% Tween and Pierce ECL western blot plus solution (Thermo Scientific, #32132) was added to the membrane for 5 minutes before imaging using the ChemiDoc XRS+ system (Bio-Rad) or ImageQuant 800 (AMERSHAM).

### RT-qPCR

Total RNA was isolated with RNeasy (Qiagen) and used to synthesize cDNA with a mixture of random hexamers and reverse transcriptase, after DNAase treatment (TAKARA PrimeScript™ RT Reagent Kit with gDNA Eraser). Diluted cDNA was used in triplicate quantitative PCR reactions with pre-tested gene specific primers and qPCRbio SYgreen Blue Mix using a QuantStudio 5 (Applied Biosystems) thermal cycler. Results were analysed using 2–ΔΔCt (relative quantitation) with quantstudio software with normalisation to housekeeping gene *Rplp0*. Statistical significance was determined using the students t-test with Holm-Sidak correction with alpha = 0.05 to identify differences in gene expression from two biological replicates using Prism.

### PyroSequencing

Genomic DNA (50-300ng) was sodium bisulfite converted using the EZ DNA Methylation-Gold Kit (#D5005 Zymo Research), eluting with 10μl of H_2_O. 1μl of bisulfite converted DNA was used as template to amplify target genomic regions using specific primers (one biotinylated) with the PyroMark PCR kit (Qiagen, #978703), following the recommended conditions (annealing temperature 56°C). 10μl of PCR reaction was used for pyrosequencing, using PyroMark Q24 advanced reagents (Qiagen, #970902) a sequencing primer, and run on a PyroMark Q24 Advanced pyrosequencer (Qiagen) with target specific dispensation orders. Statistical significance was determined using students t-test with Holm-Sidak correction with alpha=0.05 to identify differences in DNA methylation using two independent biological replicate lines, each consisting of multiple CpG site DNAme calls.

### CUT&RUN-seq

To investigate protein-DNA interactions and histone modifications locations we used the recently developed Cut and Run protocol ^57^. Dissociated single-cells were pelleted at 600xg for 3 minutes and washed twice with Wash Buffer (20mM HEPES pH 7.5, 150mM NaCl, 0.5mM Spermidine and Protease Inhibitor tablet (Roche)) at RT, then resuspended in 1ml of wash buffer. 10μl of concanavalin A-coated magnetic beads (Bangs Laboratories, #BP531), pre-washed and resuspended in Binding buffer (20mM Hepes-KOH, pH 7.9, 10mM KCl, 1mM CaCl2, 1mM MnCl2), were added to the cells and rotated for 10 minutes at RT. The bead-bound cells were isolated on a magnetic stand to remove the supernatant, and 300μl of antibody buffer (Wash buffer plus 0.02% digitonin and 2mM EDTA) with 0.5μg of antibody was added to the beads and incubated with rotation at 4°C overnight. The following day cell-bead complexes were washed with 1mL of cold Dig-Wash buffer (Wash buffer plus 0.02% digitonin) using a magnetic stand, then resuspended in 300μl of cold Dig-Wash buffer. Purified protein-A::MNase (pA-MNase) fusion was added to a final concentration of 700ng/ml and the samples were rotated for 1 hour at 4°C. Samples were washed twice in 1ml cold Dig-Wash buffer and resuspended in 50μl of Dig-Wash buffer by gentle flicking. The samples were placed in iced water to pre-cool them to 0°C. To initiate pA-MNase digestion 2μL 100mM CaCl_2_ was added and the samples were flicked to mix and returned to iced water. After 30min (histone modifications) or 5min (DPPA2) pA-MNase digestion, 50μl of 2XSTOP buffer (340 mM NaCl, 20mM EDTA, 4mM EGTA, 0.02% Digitonin, 250μg Rnase A, 250μg Glycogen) was added and the samples thoroughly mixed. Samples were incubated at 37°C for 10 minutes to release ‘CUT&RUN’ fragments from the insoluble nuclear chromatin, followed by centrifugation at 16,000xg for 5 minutes at 4°C. The supernatants were transferred to new tubes and the cell-bead complexes discarded. Subsequently, 2μl 10% SDS and 2.5μl Proteinase K was added and the samples were incubated for 10 minutes at 70°C. DNA fragments were purified and double-size selected from the suspension using SPRIselect beads (Beckman Coulter, B23318) following the manufacturers protocol for double selection (0.5x beads to DNA ratio followed by 1.3x ratio), and eluted with 30μl of 0.1xTE. CUT&RUN dsDNA samples were quantified with a Qubit III and 5-10ng used as input for library preparation using the NEBNext Ultra II DNA Library Prep Kit for Illumina (#E7645S, NEB). Libraries were prepared using the following PCR program: 98°C 30s, 98°C 10s, 65°C 10s and 65°C 5min, steps 2 and 3 repeated for 12 cycles. Library samples were paired-end sequenced on the NextSeq Illumina sequencing system (PE40).

### EM-seq

Purified genomic DNA was isolated from cells using the Zymo microprep DNA kit. To generate high quality base resolution DNA methylation libraries we used the NEBnext Enzymatic Methyl-seq (EM-seq) kit following the manufacturer’s instructions. Briefly, genomic DNA was sheared to 300bp, then end-repaired and A-tailed. Repaired DNA was then ligated to EM-seq adapters and genomic 5mC and 5hmC was oxidised to 5caC to protect methylated sites against deamination. Subsequently, APOBEC was used to deaminate unmethylated cytosines to uracils, whilst oxidized forms of 5mC/ 5hmC are not deaminated. This generates a DNA conversion system identical to bislfite conversion but with higher yields and lower duplication. The library was amplified using Q5 polymerase, and independent libraries multiplexed. These were sequenced on an Illumina Nextseq for single-end 75 (SE75).

### RNA-seq

Total RNA was collected from fresh cells using the Qiagen RNeasy kit, following the manufacturer guidelines. Total RNA was quantitated using a Quibit III and quality checked with Bioanalyser 2100 (Agilent) to ensure RIN>8.5. mRNA was enriched using the NEBNext Poly(A) mRNA magnetic isolation module and prepped into stranded libraries using the NEBnext Ultra II directional RNA library prep kit following all manufactures guidelines. Amplified libraries were multiplexed and sequenced on the NextSeq (SE80 or PE40).

### Bioinformatics analysis

#### CRISPR screen

Raw sequence reads were trimmed and quality control checked using cutadapt (v1.15) (cutadapt -g TTGTGGAAAGGACGAAACACCG) and FastQC^58^. Counting and statistical analysis of sgRNA frequency was performed with the Model-based Analysis of Genome-wide CRISPR-Cas9 Knockout (MAGeCK, version 0.5.9) tool ^59^. The normalised gRNA counts (MAGeCK -count -norm-method total) from the sorted and unsorted control samples were compared using the -test command in MAGeCK, which identifies significantly enriched/depleted gRNAs between samples. These is used to determine the relative ranking algorithm (RRA) score, which identifies enriched/depleted genes based on change in the distribution frequency of multiple independent gRNAs that target the same gene. To identify final candidates, we applied a FDR threshold of <0.05 and a fold-change (FC) gRNA frequency threshold of >3 (to select larger effect-size candidates), and intersected lists from two independent screens, each performed in independent cell lines.

#### RNA-seq

Raw reads were quality trimmed using TrimGalore (0.4.3.1, -phred33 --quality 20 --stringency 1 -e 0.1 --length 20). These were mapped to the mouse mm10 (GRCm38) genome assembly using RNA Star (2.5.2b-0, default parameters except for --outFilterMultimapNmax 1000) and reads with a MAPQ score <20 were discarded to ensure only unique-mapping high quality alignments were used for analysis of gene expression. The data was quantified using the RNA-seq quantification pipeline for directional libaries in seqmonk software to generate log_2_ reads per million (RPM) or gene-length-adjusted (RPKM) gene expression values. Where appropriate, the samples were normalised using match distribution quantitation method. Differentially expressed genes (DEG) were determined using the DESeq2 package (version, 1.24.0), inputting raw mapping counts, and applying a multiple-testing adjusted *p-value* (FDR) <0.05 significance threshold. An additional fold-change (FC) filter of >2 was applied to generate final DEGs. Differences in gene expression of the differentially methylated geneset (DMP) were tested with one-tail students t-test. For analysis of transposable elements (TE) analysis reads with a MAPQ score <20 were allowed to enable multi-mapping reads, but taking the primary alignment only. Repeat locations for mm10 (GRCm38) genome were extracted from repeatmasker, and instances overlapping or residing within 2kb of an annotated gene were excluded to prevent mixed signal derived from genic and/or TE transcription, which could confound results. Mapped reads were quantitated over these high-confidence TE, which were categorised as full-length (>5kb for LINE, >3kb for LTR) or truncated (<5kb for LINE, <3kb for LTR), enabling unbiased assessment of TE expression from WT and independent replicate mutant lines. Summed counts for all instances of each class of repeat were calculated. These were then corrected both for the total length of TE class, and the sequencing depth of individual libraries to generate log_2_ RPM expression values as previously described^60^.

#### EM-seq

Raw fastq sequences were quality− and adapter-trimmed using TrimGalore (0.4.3.1) and reads aligned to mm10 using Bismark (0.20.0), discarding the first 8 bp from the 5’ end and the last 2 bp from the 3’ of a single-end reads. Cytosine methylation status was extracted from mapped reads using the Bismark methylation extractor tool. Genome-wide methylation calls were analysed using Seqmonk software (1.44.0) with biological independent replicate datastets for each condition. To identify differentially methylated regions (DMR) the genome was first binned into sliding tiles containing 50 consecutive CpGs and their methylation status determined using the DNA methylation pipeline. DMRs were identified by running read-depth sensitive logistic regression (*p*<0.05) and binomial test (*p*<0.01) statistical filters, taking only regions scored as significant in both, with minimum of 10 reads. Differentially methylated promoters were identified by quantitating DNAme over tiles that span +/−1kb from RefSeq gene transcriptional start sites, and intersecting significant hits from both logistic regression (*p*<0.05) and binomial test (*p*<0.01) statistical filters. Differentially methylated full-length LINE1 (DML) were identified using tiles over the 5’ end of LINE1 elements >5kb (+/− 500bp) using repeatmasker annotations and intersecting logistic regression (*p*<0.05) and binomial test (*p*<0.01) statistical filters, with minimal reads of 10 per probe. To generate final DMR, DMP and DML datasets, significant hits from ESC and EpiLC *Dppa2^−/−^* were collated.

#### CUT&RUN-seq

Raw Fastq sequences were quality- and adapter- trimmed with TrimGalore (0.4.3.1, -phred33 --quality 20 --stringency 1 -e 0.1 --length 20) and aligned to the mouse mm10 genome using Bowtie2 (2.3.4.2, -I 50 -X 800 --fr -N 0 -L 22 -i ‘S,1,1.15’ --n-ceil ‘L,0,0.15’ --dpad 15 --gbar 4 --end-to-end --score-min ‘L,-0.6,-0.6’). Mapped sequences with a MAPQ score <20 were discarded. DPPA2 binding peaks were identified using MACS2 (*p*<1×10^−5^, 200nt fragments) with IgG CUT&RUN as input control. The enrichment of overlap between DPPA2 binding sites and genomic features was determined by one sample t-test against triplicate randomised genomic probesets of similar size distribution as DPPA2 binding sites. Histone modification CUT&RUN was analysed by quantifying the normalized reads over genomic features (e.g. DPPA2 binding sites, DMPs, DML, DMRs) to generate density, peak and trend datasets using seqmonk software. For analysis of repetitive elements (e.g. LINE) we allowed multi-mapping (MAPQ<20), taking only the primary alignment.

#### Gene ontology analysis

Gene ontology (GO) analyses were performed using ensemble gene ID in the DAVID (version 6.8) ^61^ tool for both differentially expressed genes (WT vs *Dppa2* KO and *Dppa4* KO in ESC and EpiLC), genes associated with differentially methylated promoters (DMPs) and screen candidates. FDR values for selected GO terms from BP_all are displayed.

---

**Supplementary Figure 1.**
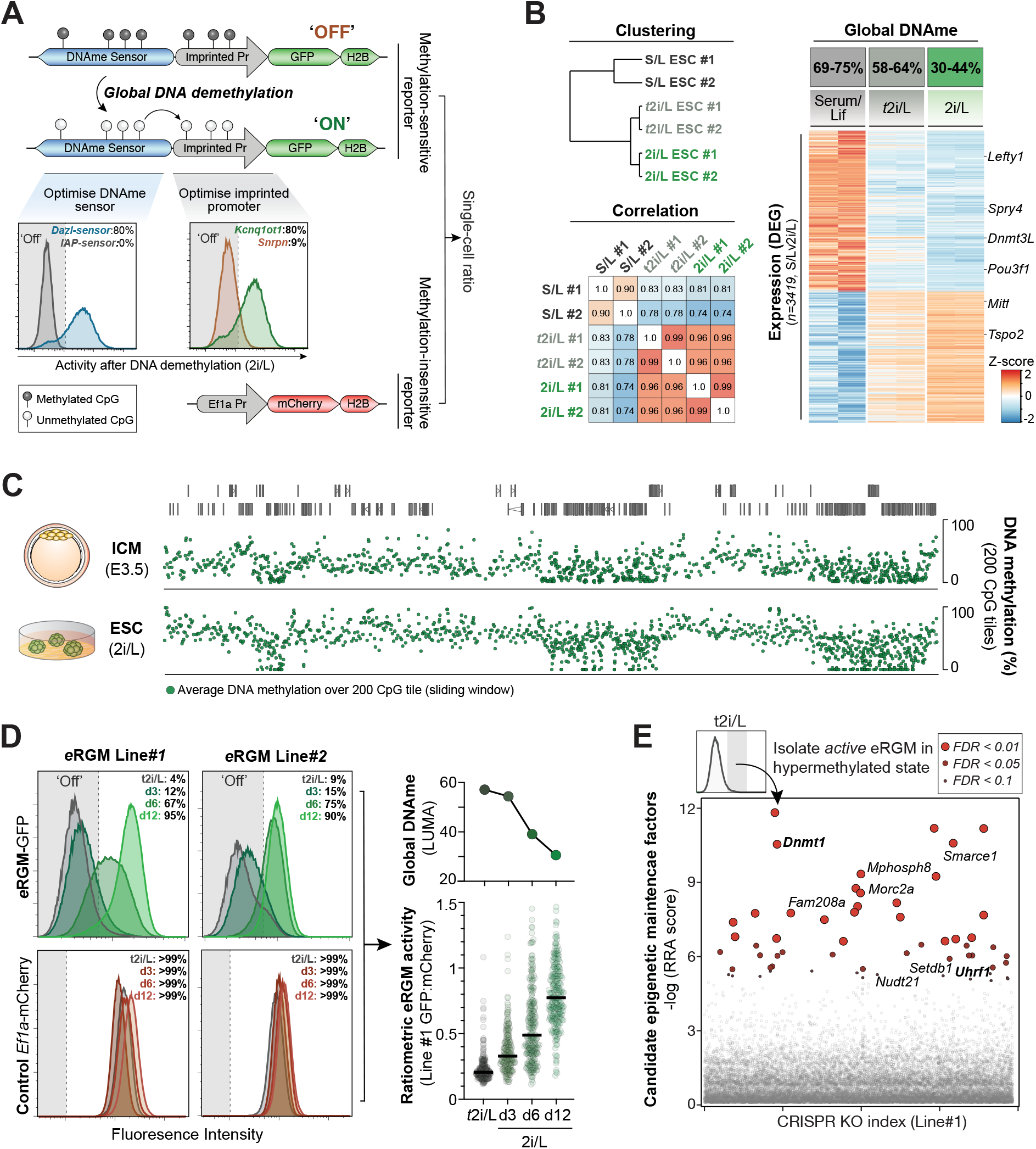
Model and enhanced ratiometric reporter for developmental DNA demethylation. **A.**) Schematic for design and optimisation of the ratiometric enhanced *e*RGM cellular DNAme reporter. The system consists of a methylation-*sensitive* imprinted promoter, which controls expression of GFP according to its level of DNA methylation. The DNA methylation level is set by an antisense upstream genomic region (DNAme sensor) that acquires a similar level of DNAme as the global DNA methylation state, and subsequently adjusts DNAme at the imprinted promoter to equivalence via proximity. This two-stage system generates a robust read-out of global DNAme status (upper panels). Changing the DNAme sensor to a region that is resistant to DNA demethylation (*e.g. IAP*) prevents eRGM activation in hypomethylated conditions (left FACS plot), confirming that the DNAme sensor status controls activity. Moreover switching the imprinted promoter from *Snrpn* to *Kcnq1ot1* enables a greater degree of expression upon DNA hypomethylation, thereby increasing the dynamic range of the reporter (right FACS plot). Shown in grey is activity in the ‘off’ hypermethyalted state. Finally, by coupling *e*RGM with a second methylation-*insensitive* reporter (*Ef1a*-mCherry), a single-cell ratiometric score can be generated that normalises for confounding factors. **B.**) Transcriptomics from ESC maintained in serum/Lif (hypermethylated), titrated *t*2i/L (hypermethylated) or 2i/L (hypomethylated). The *t*2i/L and 2i/L transcriptomes are highly comparable despite distinct global methylation states (hyper− and hypo-methylation, respectively), implying transition between these conditions isolates epigenetic resetting without confounding changes in cell identity. **C.**) Screen shot of genomic methylation pattern from naïve E3.5 epiblast and naïve ESC demonstrating *in vitro* resetting in 2i/L establishes a highly comparable methylome as *in vivo* resetting. **D.**) Representative FACS plots of progressive *e*RGM (GFP) activation by DNA demethylation during 12 day transition from *t*2i/L to 2i/L in independent *e*RGM cell lines (Line #1 and #2). Normalising to mCherry (lower panel) enables a ratiometric single-cell readout (each datapoint is a single cell), which closely tracks global methylation levels (right panel). **E**.) CRISPR screen for gene KO that enable *e*RGM activation even under hypermethylated conditions (*t*2i/L) identifies key known DNA methylation and chromatin regulators, confirming *e*RGM specificity and sensitivity to modulation of epigenetic systems.

**Supplementary Figure 2.**
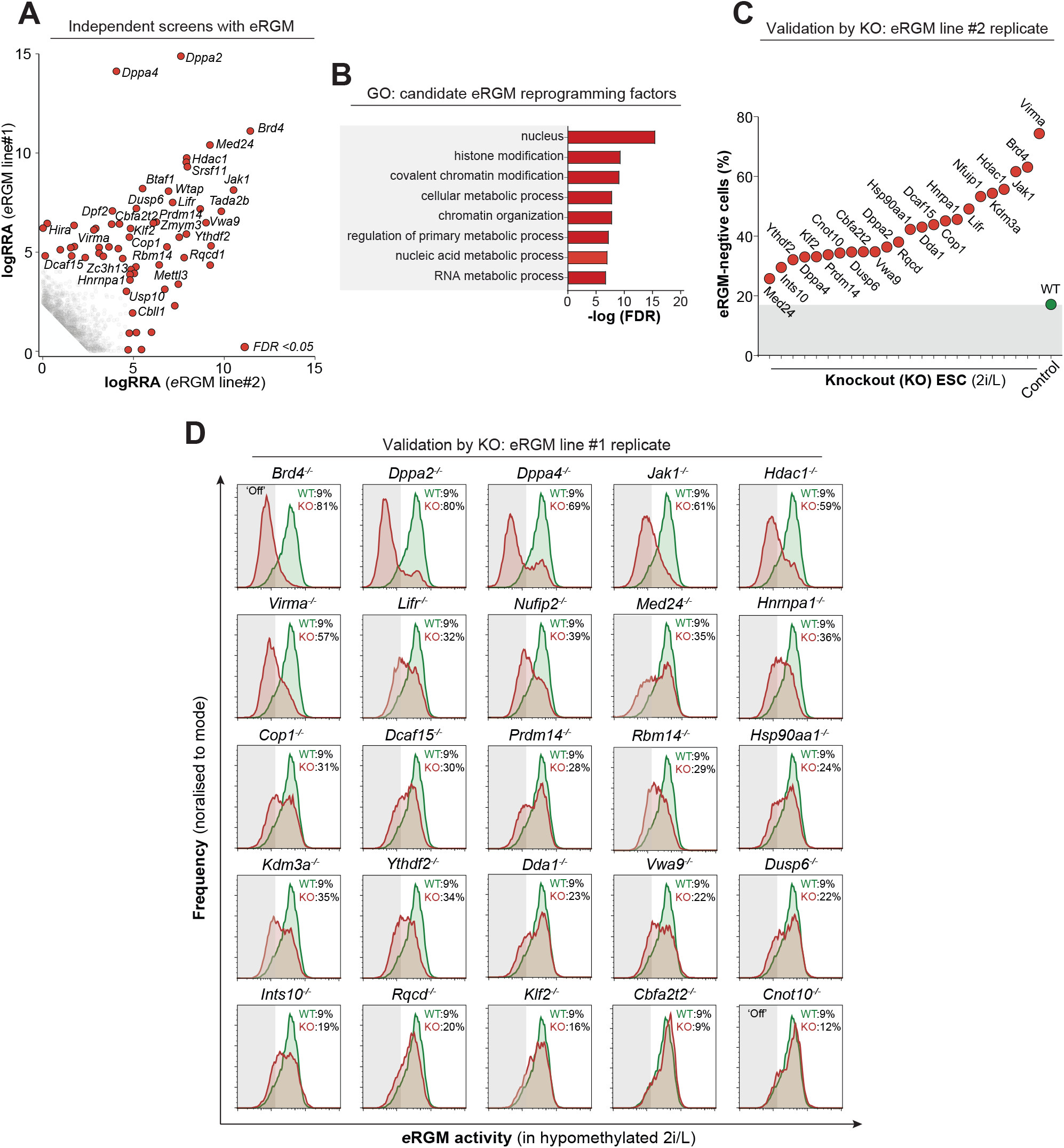
Validation of CRISPR screen candidates for epigenetic reprogramming. **A.**) Scatter plot showing significance (RRA) values for candidates from independent screens of independent cell lines are highly correlated. *FDR*<0.05 was used as a threshold. **B.**) Gene ontology (GO) analysis of candidate factors from the screen shows enrichment for nuclear activity, and involvement in chromatin and/or nucleic acid processes consistent with being epigenetic regulators. **C.**) Percentage of cells that remain *e*RGM-negative in hypomethylation-inducing 2i/L upon knockout of the indicated candidate reprogramming factor. Shown is data from KO generated independently in *e*RGM line#2, analogous to Fig 2A in *e*RGM line#1 **D.**) Representative flow cytometry histograms demonstrating knockout of most candidates leads to a significant block of *e*RGM activation amongst single cells, implying altered epigenetic resetting. Shown is percentage single-cells with *e*RGM-negative ‘off’ (marked in grey) after 12 days in DNA hypomethylation-inducing 2i/L culture.

**Supplementary Figure 3.**
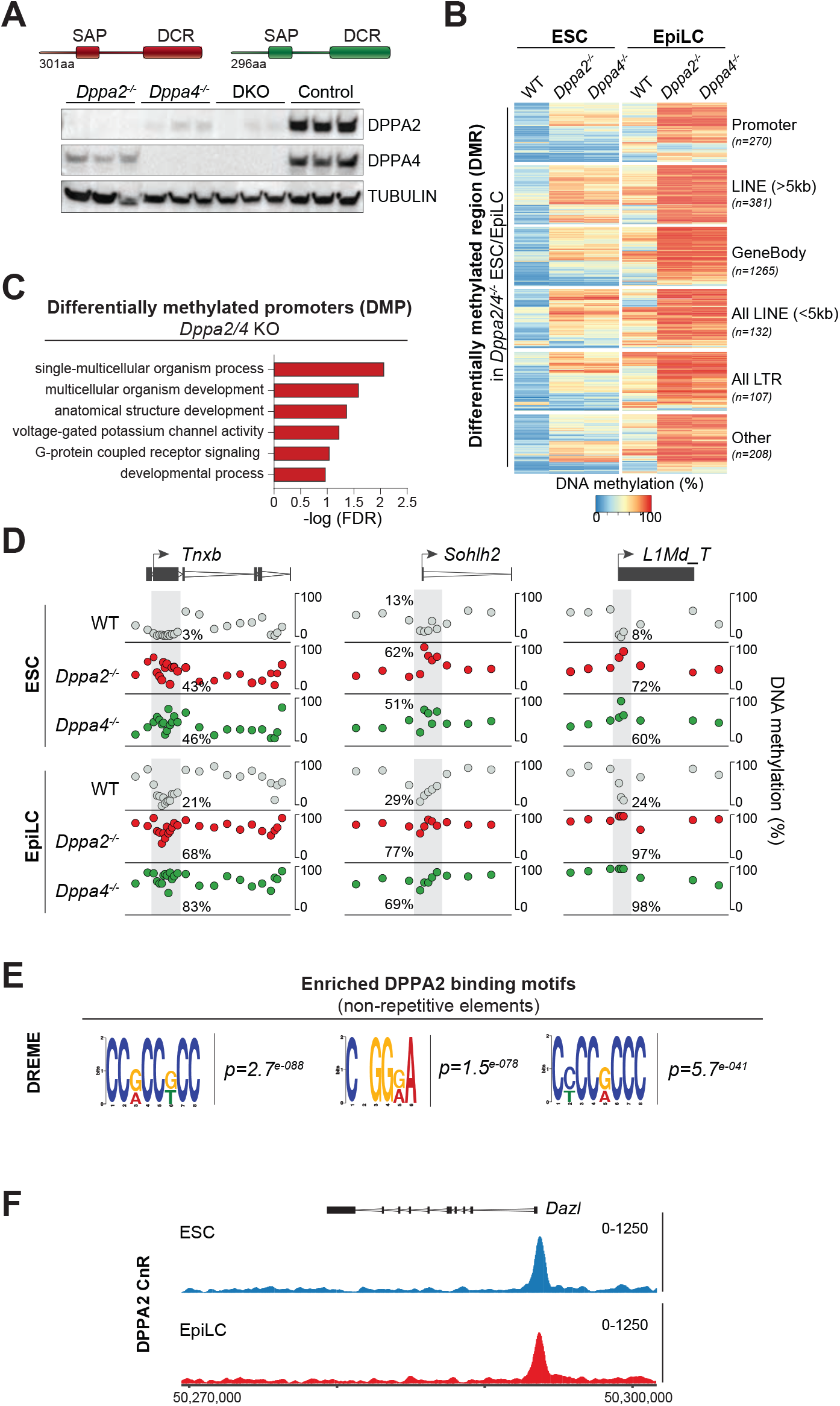
The DNA methylation landscape in *Dppa2/4* knockout. **A.**) Western blot confirming loss of DPPA2, DPPA4, or both (DKO) protein(s) upon generation of clonal knockout ESC lines. Note loss of DPPA2 protein leads to depletion of DPPA4, and reciprocally, potentially due to disruption of the heterodimeric complex that stabilises each protein. **B.**) Heatmap showing methylation status of significant differentially methylated regions (from sliding 50 CpG windows) (DMR) in *Dppa2* knockout cells. Note *Dppa4* DMRs are highly correlated with *Dppa2*. **C.**) Gene ontology (GO) of genes associated with differentially methylated promoters (DMP) in *Dppa2* KO ESC or EpiLC reveals enrichment for developmental-associated gene classes. **D.**) Genome view showing DNA methylation patterns in WT and *Dppa2/4* KO ESC and EpiLC. Each datapoint represents the windowed average methylation of 20 CpG sites. **E.**) Top three enriched DPPA2 binding motifs reveal preference for GC. **F**.) Genome view showing that DPPA2 binds at the genomic sensor region used in eRGM (*Dazl)* in ESC and EpiLC and protects it from *de novo* methylation.

**Supplementary Figure 4.**
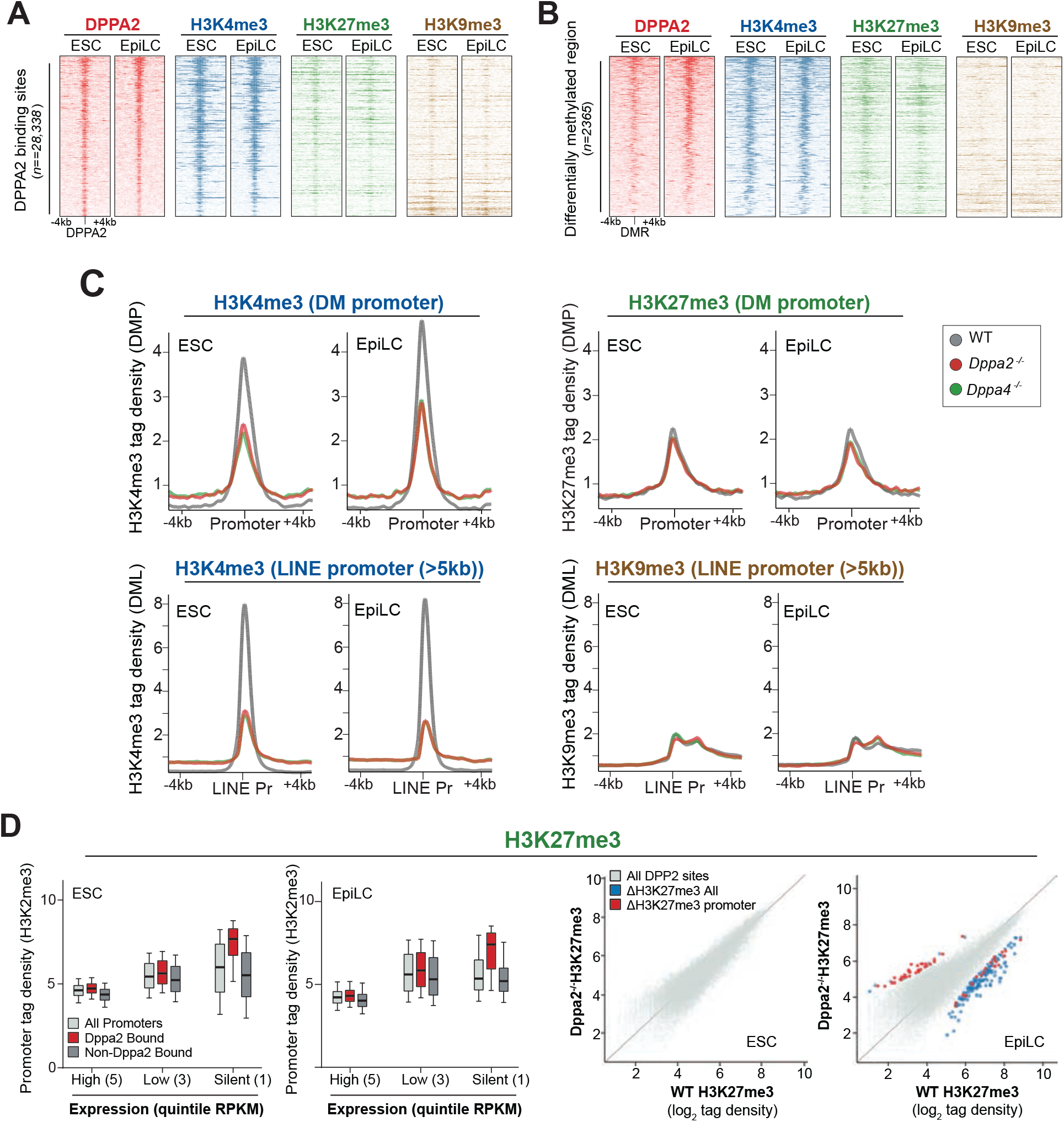
Chromatin state at DPPA2 binding sites and upon knockout. **A.)** Aligned probe plot showing enrichment of H3K4me3, H3K27me, and H3K9me3 centered on DPPA2 binding sites with 4kb either side in WT ESC. Plots are ordered equivalently, by DPPA2 binding enrichment, which is shown in red. H3K4me3 shows strong enrichment over nearly all DPPA2 sites, whilst H3K27me3 shows modest enrichment. **B.)** As in A, but chromatin enrichment is centered on differentially methylated regions (DMR). H3K4me3 is enriched in WT ESC at sites prone to hypermethylation upon *Dppa2/4* knockout **C.)** Trend plot showing the enrichment of chromatin marks over gene promoters (upper panels) or over full-length LINE1 promoters (lower panels) that acquire DNAme in *Dppa2 KO*. H3K4me3 exhibits a dramatic depletion in *Dppa2* or *Dppa4 KO*, whilst H3K27me3 and H3K9me3 exhibit no significant changes at these loci. **D.)** Left: Boxplot showing H3K27me3 enrichment over DPPA2-bound promoters binned for expression quintile. Right: Scatter plot showing that, unlike H3K4me3, very few DPPA2 binding sites exhibit a change in H3K27me3 upon *Dppa2* knockout.

**Supplementary Figure 5.**
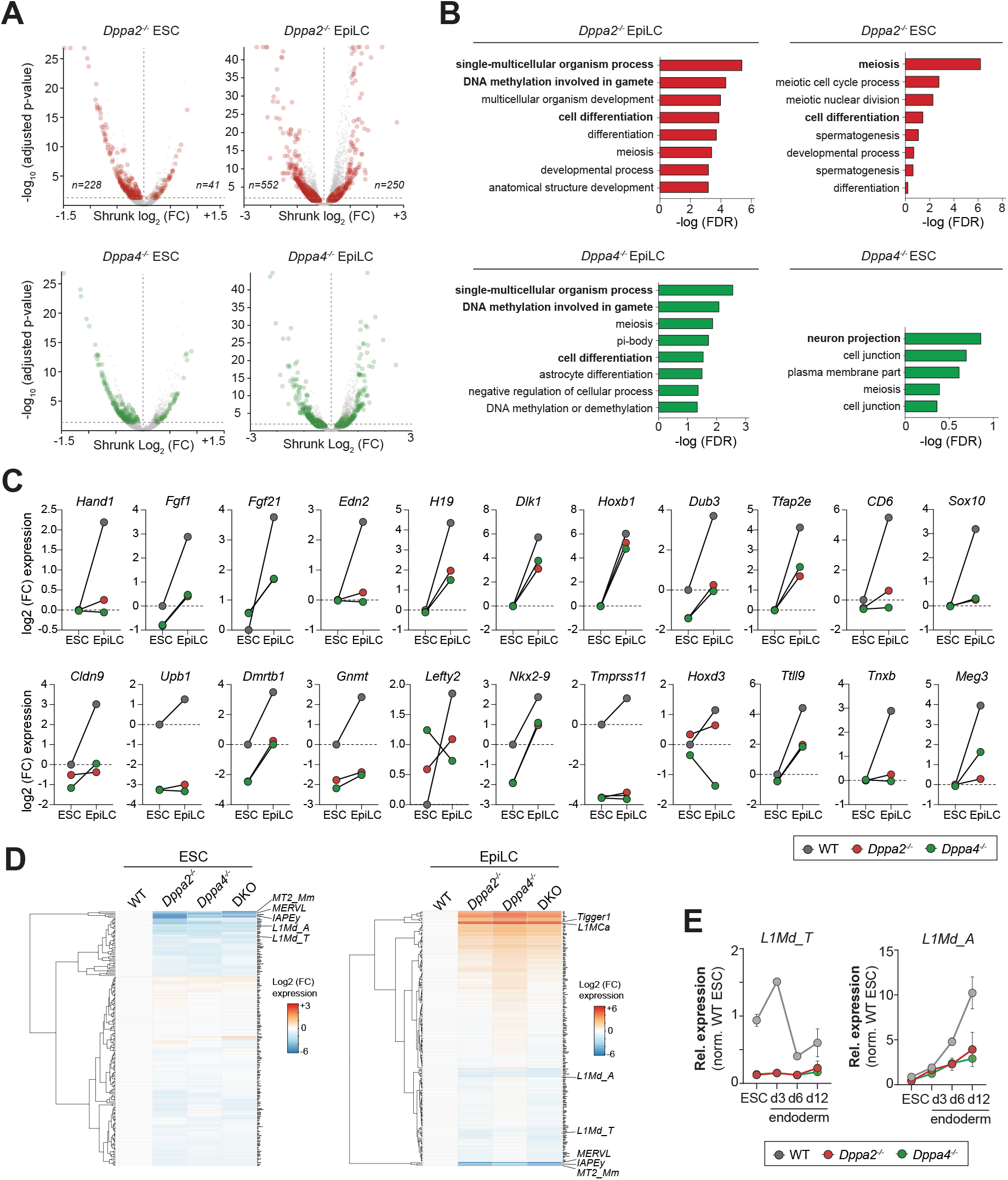
Transcriptional and developmental competence upon Dppa2/4 deletion. **A.)** Volcano plot showing all differentially expressed genes (DEG) from *Dppa2/4* KO ESC and EpiLC. Note many more genes are repressed than activated. **B.)** Gene ontology (GO) of DEGs in ESC and EpiLC reveals a strong enrichment for developmental-associated terms, which is driven by silenced developmental genes in *Dppa2/4* KO **C.)** Representative examples of developmental genes that fail to active in *Dppa2/4* KO EpiLC. **D.)** Heatmap showing expression of all transposable elements (TE) in mutant cells. **E.)** qRT-PCR confirming that evolutionary young *L1Md_T* and *L1Md_A* exhibit impaired expression in *Dppa2/4* KO, implying *Dppa2/4* are required to maintain competence for TE activity.

**Supplementary Figure 6.**
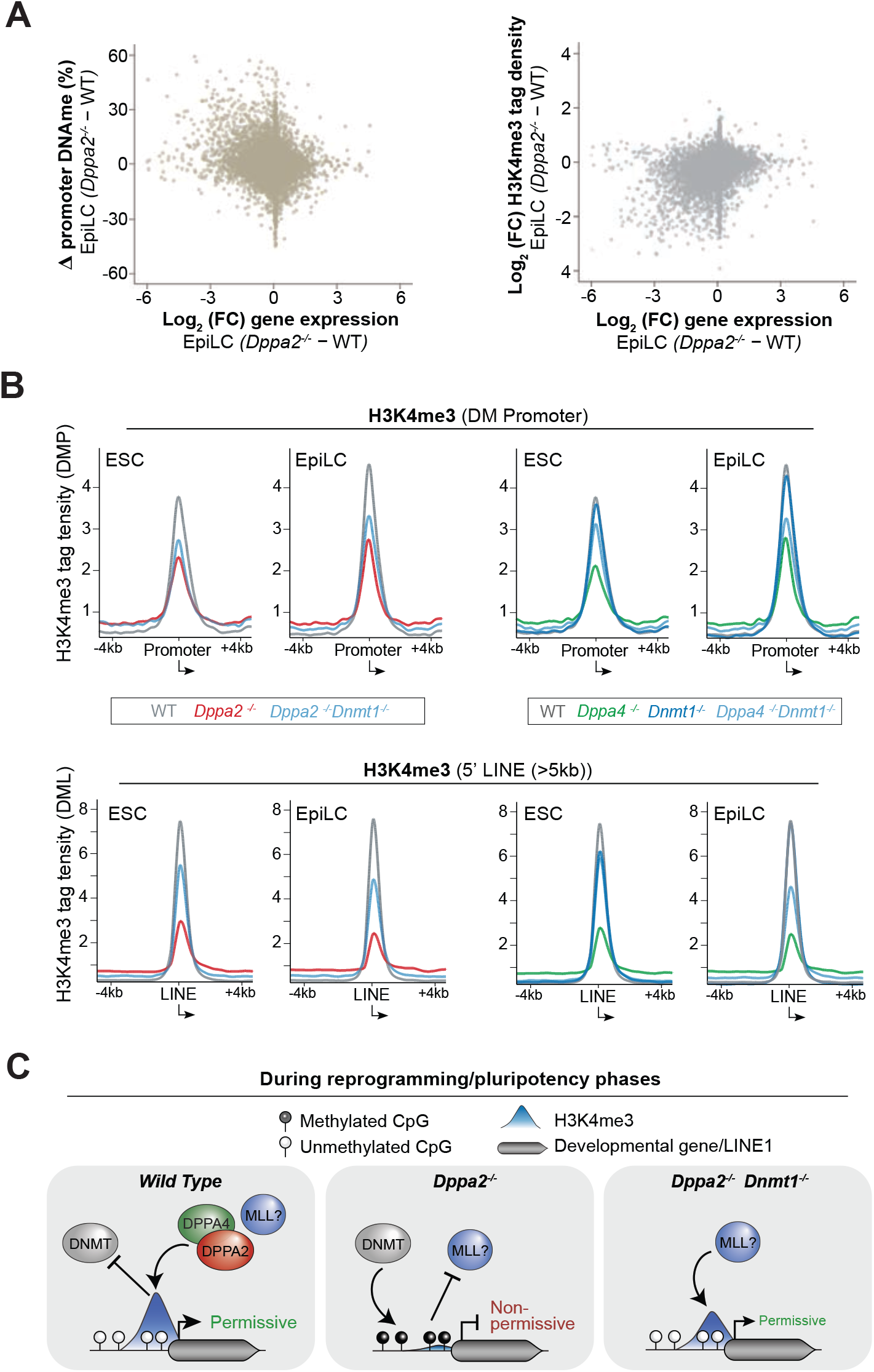
Functional interaction between DNAme, H3K4me3, and gene silencing. **A.)** Scatter plot showing inter-relationships between changes in H3K4me3 and DNA methylation versus gene expression upon *Dppa2 KO* in EpiLC **B.)** Trend plots showing the enrichment of H3K4me3 over gene promoters (upper panels) and over full-length LINE1 promoters (lower panels) that acquire DNAme in *Dppa2 KO*. Shown for WT, *Dppa2 KO* (or *Dppa4 KO*), *Dnmt1 KO* and *Dnmt1/Dppa2 KO* (*or Dppa4 KO*) in ESC and EpiLC**. C.)** Proposed model of the interplay between H3K4me3 and DNA methylation at *Dppa2/4* targets in regulating gene expression competence.

